# Expanded palette of RNA base editors for comprehensive RBP-RNA interactome studies

**DOI:** 10.1101/2023.09.25.558915

**Authors:** Hugo C. Medina-Munoz, Eric Kofman, Pratibha Jagannatha, Evan A. Boyle, Tao Yu, Krysten L. Jones, Jasmine R. Mueller, Grace D. Lykins, Andrew T. Doudna, Samuel S. Park, Steven M. Blue, Brodie L. Ranzau, Rahul M. Kohli, Alexis C. Komor, Gene W. Yeo

## Abstract

RNA binding proteins (RBPs) are key regulators of RNA processing and cellular function. Technologies to discover RNA targets of RBPs such as TRIBE (targets of RNA binding proteins identified by editing) and STAMP (surveying targets by APOBEC1 mediated profiling) utilize fusions of RNA base-editors (rBEs) to RBPs to circumvent the limitations of immunoprecipitation (CLIP)-based methods that require enzymatic digestion and large amounts of input material. To broaden the repertoire of rBEs suitable for editing-based RBP-RNA interaction studies, we have devised experimental and computational assays in a framework called PRINTER (<u>p</u>rotein-<u>R</u>NA <u>in</u>teraction-based triaging of <u>e</u>nzymes that edit <u>R</u>NA) to assess over thirty A-to-I and C-to-U rBEs, allowing us to identify rBEs that expand the characterization of binding patterns for both sequence-specific and broad-binding RBPs. We also propose specific rBEs suitable for dual-RBP applications. We show that the choice between single or multiple rBEs to fuse with a given RBP or pair of RBPs hinges on the editing biases of the rBEs and the binding preferences of the RBPs themselves. We believe our study streamlines and enhances the selection of rBEs for the next generation of RBP-RNA target discovery.

## Introduction

RNA-binding proteins (RBPs) bind to RNA regulatory elements to regulate the RNA lifecycle of networks of RNA species. As disruption of protein-RNA interactions is associated with many human diseases^1^, scalable technologies that identify protein-RNA interactions are critically needed to provide deeper insights into RNA regulation. Immunoprecipitation (IP)-based strategies coupled with high-throughput sequencing such as cross-linking immunoprecipitation (CLIP) are routinely used to identify RBP targets and binding sites of RBPs. The eukaryotic ribosome is also composed of a complex of RBPs, and ribosome profiling methods such as Ribo-seq^2^ and variants that leverage enhanced CLIP^3^ with antibodies recognizing ribosome subunit proteins can evaluate the mRNA translatome^3^. Unfortunately, these techniques rely on the digestion of the unprotected portions of the interacting RNA, hampering the discrimination of RBP binding sites on alternative mRNA isoforms and the association of multiple proteins with the same transcript.

Technologies such as TRIBE^4^ and STAMP^5^ address these issues by fusing RNA base editors (rBEs) to full-length RBPs, thereby obviating the need for RNase digestion and protein-RNA cross-linking. In STAMP, RBP-APOBEC1 fusions yield statistically significant clusters of C-to-U edits near the known RBP binding motif^5,6^. Since RNase digestions are nonobligatory in STAMP, long-read sequencing detected RBP interactions with specific mRNA isoforms^5^. STAMP requires fewer cells than CLIP and was demonstrated to enable single-cell analysis of RBP-RNA interactions^5^. However, the enzymes used in TRIBE and STAMP have reported native sequence context preferences for base deamination. APOBEC1 prefers editing cytosines that are flanked by adenosine (A) and uridine (U) bases^7^ and disfavors editing sites with upstream guanosine^8^. The catalytic domains of TRIBE enzymes (ADARcd and the mutated derivative used for hyperTRIBE) have a strong preference for editing double-stranded RNA, even without the double-stranded RNA-binding domains, and this leads to false negatives in TRIBE experiments^4,9–14^. Hence, we contend that the current paucity of available rBEs constitutes a constraint in the pursuit of transcriptome-scale exploration of protein-RNA interactions.

To substantially expand the repertoire of rBEs, we developed an experimental and computational framework consisting of a combination of reporter constructs and transcriptome-wide analysis in live human cells we term PRINTER (<u>p</u>rotein-<u>R</u>NA <u>in</u>teraction-based triaging of <u>e</u>nzymes that edit <u>R</u>NA) that evaluated the editing activities and specificities of 31 rBEs. We evaluated the most promising rBE candidates through their fusion with two distinct full-length human RBPs, RBFOX2 and RPS2, each known for their unique RNA interaction preferences. Our experiments successfully identified seven new enzymes capable of detecting transcriptome-wide protein-RNA interactions with high sensitivity and specificity. This expansion builds upon the foundation established by the previous three enzymes^4,5,15^, significantly broadening the toolkit for RNA interaction studies. Furthermore, our comprehensive characterization of editing biases associated with different rBEs when fused to RBPs underscores the importance of considering these biases, especially when selecting single or multiple rBE fusions. This choice becomes particularly crucial when studying RBPs with strict sequence motif preferences, such as RBFOX2, in contrast to RBPs with more broad sequence specificity, such as RPS2. Lastly, we recommend pairs of rBEs that are well-balanced for enabling dual-RBP editing measurements on the same RNA transcript. Our study sets the stage for enabling the next stage of discoveries that leverage diverse rBEs to illuminate RBP and ribosome-RNA interactions transcriptome-wide.

## Results

### A two-component reporter system evaluates rBEs for RNA editing activity in cells

To identify ideal fusion partners to full-length RBPs, we designed a framework to evaluate the activities of putative RNA base editors (rBEs) in live mammalian cells. Our system comprises two components that are transiently co-transfected into human embryonic kidney (HEK293XT) cells. The first is an editable mRNA reporter that consists of a 3’UTR with twelve MS2 bacteriophage stem-loops (12X MS2-SLs) that physically interact with the RBP MS2 bacteriophage coat protein (MCP)^16^. The second component is an MCP-rBE fusion whose expression is controlled by doxycycline 24 hours after transfection (Figure 1A, B). Recruitment of the rBE to the reporter by the MCP interaction with MS2-SLs results in RNA editing on nearby substrate bases (Figure 1B). After 48 hours, DNA-free RNA is isolated from lysed cells and targeted RNA sequencing then selectively detects edits on the reporter mRNA (Supplementary Figure 1A). As a positive control, we evaluated MCP-APOBEC1 used previously in STAMP^5^. To ensure the robustness of the results, two biological replicates were conducted for the experiments. As expected, APOBEC1 deposited C-to-U edits on the reporter mRNA. Sequencing identified multiple instances of uridine (U) in place of cytosine (C) at several positions along the twelve-MS2 stem-loop region, indicating C-to-U editing by the MCP-APOBEC1 fusion at those sites (Figure 1C, green bars). In contrast, we only observed low levels of edits that may be due to expected PCR or sequencing errors in the absence of the MCP-APOBEC1 fusion (“No rBE,” Figure 1C). Therefore, our two-component system successfully detects rBE-mediated RNA editing.

**Figure 1:**
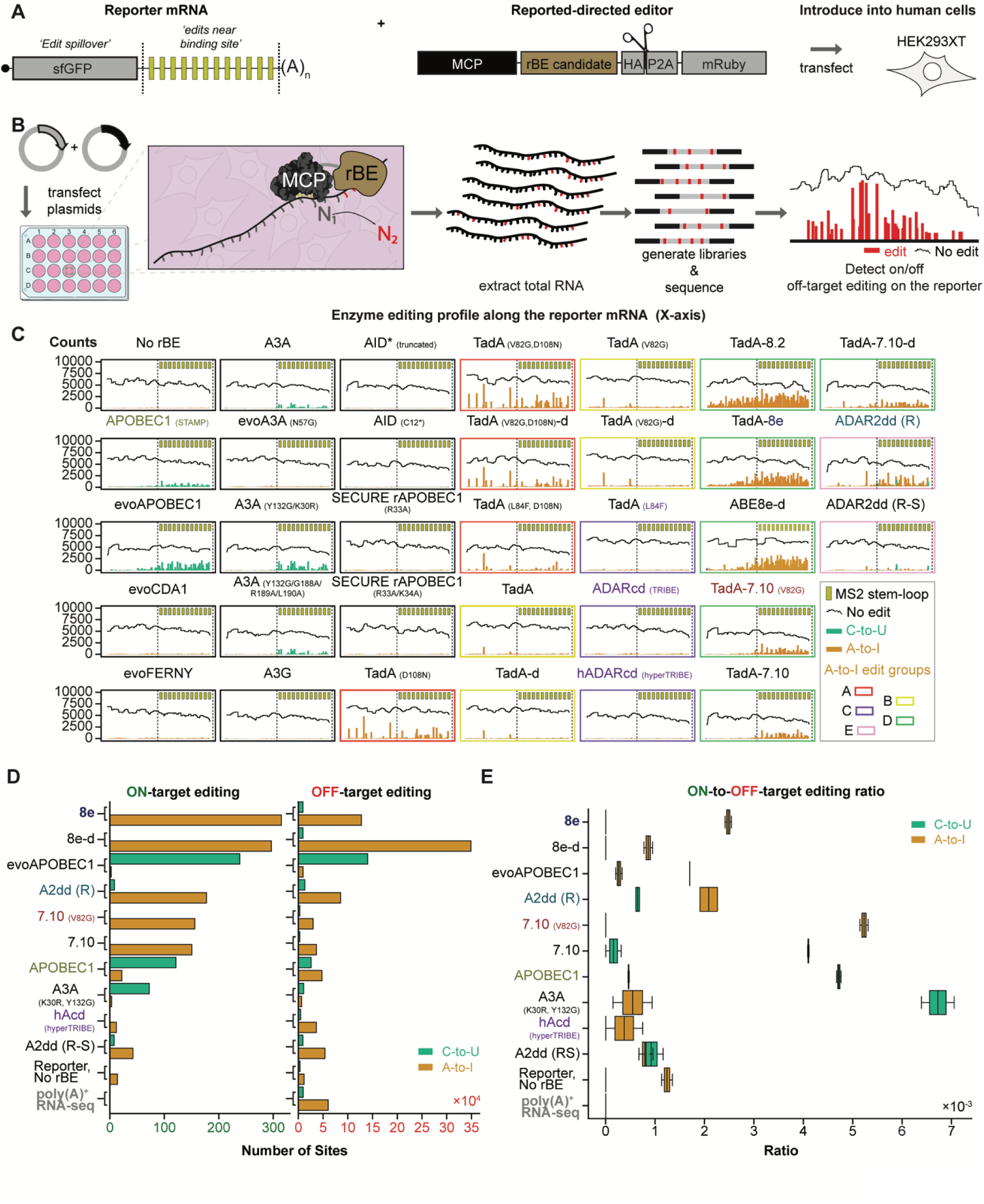
A reporter system to assay RNA base editing activity of novel rBEs. **A)** Reporter mRNA bearing twelve MS2 bacteriophage stem-loops (yellow bars) downstream of a Super folder GFP coding sequence (sfGFP) and a chimera bearing an MS2 coat-protein (MCP, black) domain fused to a candidate RNA base-editing enzyme (rBE, brown). **B)** Strategy to test candidate rBEs. Plasmids encoding the constructs in A) are co-transfected into HEK293XT cells so that the MCP binds the MS2 stem-loops in the reporter, and the rBE catalyzes RNA editing. After total RNA is isolated, targeted RNA sequencing is used to detect edits along the reporter sequence. **C)** Fraction of total covered bases at each position along the twelve MS2 stem-loop reporters exhibiting either C-to-U (green), A-to-I (orange), or no (black horizontal line) edits. **D)** The number of C-to-U (green) and A-to-I (orange) edits on the reporter mRNA (on-target, left) or other poly(A)+ RNAs (off-target, right). **E)** Ratio of the number of on-target C-to-U (green) and A-to-I (orange) edits on MS2 twelve loop construct vs. the number of off-target edits on poly(A)+ RNAs, plotted for replicates across each enzyme.

### Transcriptome-wide analysis reveals rBEs display a range of editing activity and accuracy

We evaluated rBEs that previously demonstrated precise RNA editing (e.g., RESCUE-S ADAR2dd^17^, hereafter A2dd (R-S)) and DNA base editors with high levels of “on-target” DNA editing activity (e.g., evoCDA1^8^) or “off-target” RNA editing activities (e.g., TadA-7.10 (V82G), hereafter 7.10 (V82G)) from previous studies that targeted editing to specific DNA or RNA loci using or Cas9- or Cas13-based technology^8,17–29^. We evaluated 31 candidate editors, including fourteen C-to-U editors (including APOBEC1), seventeen A-to-I editors (including TRIBE and hyperTRIBE enzymes), and two capable of catalyzing both A-to-I and C-to-U edits. Our panel of enzymes includes different protein families such as *Escherichia coli* (*E. coli*) tRNA-specific adenosine deaminase (TadA), Activation-induced cytidine deaminase (AID)/Apolipoprotein B mRNA editing enzyme, catalytic polypeptide-like (APOBEC), and adenosine deaminase acting on RNA (ADAR). These enzymes were expressed as C-terminal fusions to MCP (Figure 1A, B) and individually evaluated by our reporter system in two biological replicates each.

The editors deposited a wide variety of C-to-U editing profiles on the reporter RNA. The mutant derivative of APOBEC1, evoAPOBEC1^8^, yielded noticeably higher editing near the MS2 stem-loops than APOBEC1, albeit with edits reaching farther upstream into the GFP coding sequence (“edit spillover,” Figure 1C). Since the GFP coding sequence is away from the MCP-binding sites, editing there may reflect off-target RNA editing (e.g., editing while the MCP is not bound to the RNA). The APOBEC3A-based mutant, A3A (Y132G/K30R)^30^, similarly catalyzed higher editing activity near the MCP binding sites than APOBEC1. However, A3A (Y132G/ K30R) had far less edit spillover, which may be a desirable feature for editing-based identification of binding sites of sequence-specific RBPs (Figure 1C). Furthermore, the wild-type A3A and the A3A (Y132G/G188A/R189A/L190A)^30^ mutant produced similar editing rates but different editing patterns than APOBEC1, indicating that these enzymes may complement APOBEC1 (Figure 1C). Lastly, the C-to-T-editing enzymes evoCDA1^8^, evoFERNY^8^, evoA3A(N57G)^18^, APOBEC3G (A3G)^31^, AID* (truncated)^32^, AID (C12*)^33^, SECURE rAPOBEC1 (R33A)^18^, and SECURE rAPOBEC1 (R33A/K34A)^18^ did not produce detectable RNA editing on our reporter mRNA (Figure 1C, see also Materials and Methods), confirming the engineered reduction of RNA-editing for the SECURE rAPOBEC1 DNA-editing enzymes^18^.

The A-to-I RNA-editing enzymes we assessed produced at least five editing patterns on the reporter RNA. The first group of editors, group A, deposited edits across the entirety of the reporter mRNA (TadA (D108N)^21^, TadA (V82G/D108N)^21^, TadA (V82G/D108N)-d^21^, and TadA (L84F/D108N)^21^, Figure 1C). Group B editors interestingly primarily edited the GFP coding region with little to no editing near the 3’UTR MCP binding sites (TadA^21^, TadA-d^21^, TadA (V82G)^18,21^, TadA (V82G)-d^18,21^), while Group C editors barely had any signal which we suspect arose from PCR or sequencing errors (TadA (L84F)^21^, ADARcd^4^, and ADARcd (E488Q)^15^ (hereafter hAcd (hyperTRIBE) for simplicity) (Figure 1C). Group D editors primarily edited near the MS2 binding sites, albeit along a wide range of spillover rates (editing observed on the GFP coding sequence) with enzymes such as TadA-8.2^25^ TadA-8e^25^, and TadA-8e-d^25^ (hereafter 8.2, 8e, and 8e-d respectively) at the high end and enzymes like 7.10 (V82G)^18^, TadA-7.10^18,21,34^ (hereafter 7.10), and TadA-7.10-d^18,21,34^ (hereafter 7.10-d) at the low end (Figure 1C). The last group of editors, group E, comprised ADAR-derivatives RESCUE ADAR2dd^17^ (hereafter A2dd (R)) and A2dd (R-S)^17^, edited both A-to-I and C-to-U simultaneously, an activity also observed with Cas13-directed RNA editing^17^. However, despite similar editing capability and spillover rates, the A2dd (R) enzyme displayed much more activity than the A2dd (R-S) mutant (Figure 1C). We demonstrate that our reporter system successfully evaluated A-to-I and C-to-U RNA editing activity and specificity.

In our reporter assays, we pinpointed eight enzymes that outperformed TRIBE and STAMP in terms of signal-to-noise ratio. Among these, A3A (Y132G/K30R), 7.10 (V82G), and 7.10 displayed the most potent on-target editing within the reporter mRNA’s 3’UTR, while maintaining minimal off-target edits on the GFP coding region (see Figure 1C). EvoAPOBEC1, 8e, 8.2, 8e-d, and 8.2-d exhibited remarkable editing activity in proximity to the MCP binding sites. These enzymes are particularly valuable for studying proteins with brief dwell times on their target RNAs, despite some detectable spillover rates (refer to Figure 1C). Importantly, the spillover appears to be primarily associated with the robust interaction between MCP and the cognate MS2 stem-loops (as indicated by a low dissociation constant, K_d_ = 10^-9^–10^-8^ M (ref. ^35^). Hence, we hypothesize that RBPs with shorter dwell times on RNA will yield more precise editing near genuine RBP binding sites, with manageable off-target effects. Additionally, we included A2dd (R) and A2dd (R-S) due to their capacity to edit both adenosine and cytidine, expanding the range of sequence substrates that can be captured.

### Detecting genome-wide off-target editing for the top editing candidates

To determine the editing accuracy for the seven candidates compared to TRIBE and STAMP, we reasoned that since the reporter bears sequences that are not native to HEK293XT cells (namely the GFP and the MS2 SLs), we can gauge the accuracy of the MCP-rBE candidate by comparing the frequency at which the fusion edits the reporter (on-target editing) to the edits on endogenous transcripts that lack the MCP binding sites (off-target editing, Figure 1D). We performed polyA+ RNA sequencing on the same RNA samples from both experimental replicates used to prepare the targeted RNA sequencing in the reporter-based screen (Figures 1A, B, C, and D). Analysis of transcriptome-wide data revealed strong agreement with our targeted RNA-seq assay results, uncovering the highest editing rates for 8.e and 8.e-d, then decreasing progressively in editing rates to evoAPOBEC1, A2dd (R), 7.10 (V82G), 7.10, APOBEC1, A3A (Y132G/ K30R) and A2dd (R-S) (Figure 1D). The editing rate for hAcd (hyperTRIBE) enzyme was comparable to or slightly below background editing (Figure 1D, E). Further, the off-target editing rates for each enzyme generally correlated with editing activity, with the most active editors (e.g., 8e) demonstrating higher off-target editing than the least active editors (e.g., A2dd (R-S)). Intriguingly, the 8e-d di-mer demonstrated lower on-target editing activity while simultaneously increasing off-target editing relative to the 8e monomer (Figure 1D, E). Our results indicate that rBEs 7.10 (V82G) and A3A (Y132G/K30R) have the highest editing and signal-to-noise activity of the A-to-I and C-to-U rBEs, respectively.

### RBFOX2 fusions with candidate rBEs reveal distinct but authentic RNA targets

Having established that our enzymes performed well as fusions to MCP, we next assessed whether our candidate rBEs demonstrate accurate editing when fused to distinct human RBPs. We engineered 7.10 (V82G), A2dd (R), APOBEC1, and 8e ORFs as C-terminal fusions of RNA Binding Fox-1 Homolog 2 (RBFOX2) protein (Figure 2A, B). We reasoned that since RBFOX2 binds specific sequence motifs ((U)GCAUG^36^, where (U) is present in some, but not all binding sites), fusing the candidate rBEs to this RNA-binding protein should yield edits near the known RBFOX2 binding motif. To distinguish RBFOX2-directed editing from free rBE -directed editing, we also expressed each rBE without fusion to an RNA-binding protein domain (“enzyme-only”; Figure 2A, B). In biological triplicates, we transiently transfected HEK293XT cells with plasmids encoding each fusion. After 72 hrs of inducing expression of the constructs, we generated and sequenced RNA-seq libraries. The sequencing data for RBP-fusions and enzyme-only experiments was first processed with the SAILOR algorithm to detect both A-to-I and C-to-U base changes^5,37^, before we used the FLARE algorithm^6^ to identify statistically significantly enriched edit clusters along the exons and introns of the target RNA species (Figure 2B). We pinpointed clusters consistently identified in all three RBFOX2-rBE experiments and designated them as RBFOX2-rBE edits. Conversely, clusters present in all three enzyme-only experiments were categorized as background noise and subsequently excluded from the RBFOX2-rBE experimental dataset. We retained 7,882 clusters (representing 4,151 genes) for 8e, 4,003 (2,437 genes) for A2dd (R), 1,897 (1,274 genes) for APOBEC1, and 736 (549 genes) for 7.10 (V82G) (Figure 2C, D). Our transcriptome-wide results were generally consistent with the reporter assay findings, as the enzymes with the highest number of clusters (8e, A2dd (R)) also achieved the greatest editing on the reporter (Figure 2C, D, and Figure 1D). The only inconsistent observation was with APOBEC1, which when fused to RBFOX2, resulted in more clusters and identified more targets than 7.10 (V82G), but it produced less editing on the reporter than 7.10 (V82G) (Figure 2C, D, and Figure 1D). This may be due to enzyme-specific biases, although it may be that the differences are within margins of error. Despite the differences in the number of clusters detected by each fusion, de novo motif enrichment analyses using HOMER^38^ identified statistically significant enrichment of the central GCAUG motif for all RBFOX2-rBEs evaluated (Figure 2E, and Supplementary Figure 2A, B, and C). We observed that each of the RBFOX2-rBE fusions deposited edits near the GCAUG sequence statistically (p-value <0.00001) more frequently than enzymes only (Figure 2F, G). It is worth noting that the proportion of edit clusters containing the core RBFOX2 binding motif (GCAUG) may appear relatively low across various enzymes. However, these findings align with independent CLIP studies, revealing that in HepG2 cells and HEK293, less than 33% and approximately 40% of RBFOX2 binding sites, respectively, feature the typical RBFOX2 binding motif^39,40^. In addition, many edited regions lacking the canonical motif may contain RBFOX2 motifs of intermediate affinity (see ref ^6^). More importantly, empirical permutation tests demonstrated that the RBFOX2-rBE clusters were more likely to coincide with RBFOX2-APOBEC1 eCLIP peak sequences than randomly selected, similarly sized regions (Figure 2H). *A priori,* we do not expect a perfect overlap between rBE- and eCLIP-based detection of RBFOX2 binding sites (Figure 2H).

**Figure 2:**
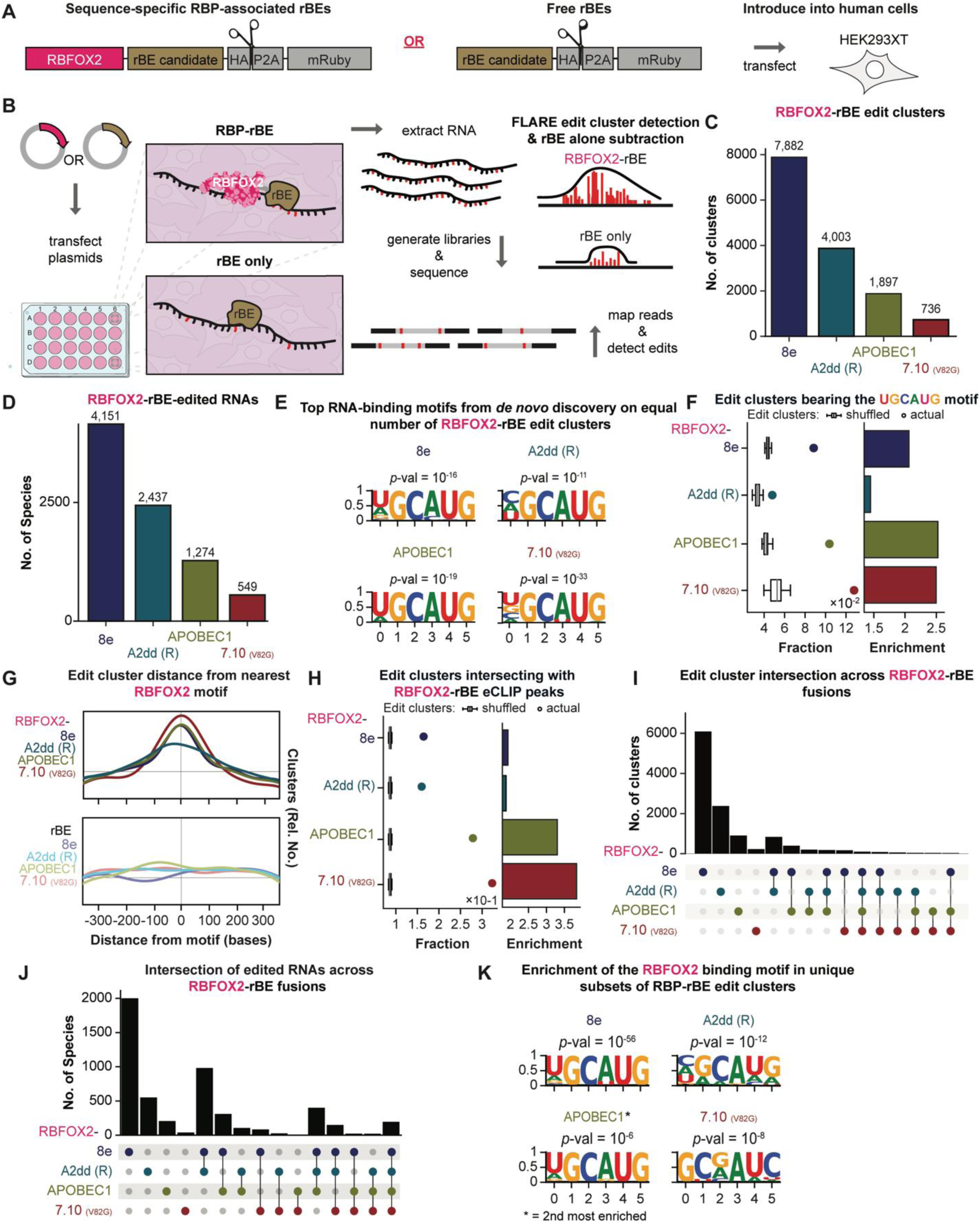
RBP-mediated RNA editing with RBFOX2 and the top rBE candidates. **A)** Constructs bearing either the RBFOX2 RNA-binding protein (pink) fused to a candidate rBE (brown) at its C-terminal end (“Sequence-specific RBP-associated rBEs,” left) or a candidate rBE (brown) without an RBP domain (“Free editors,” right). **B)** Strategy to detect RBFOX-2 directed or free rBE activity. Plasmids encoding each construct are separately transfected into HEK293XT cells. In the cell, the rBEs fused to the RBFOX2 RBP are predicted to edit near the cognate GCAUG core binding site, while the free enzyme is not. After 72 hrs, total RNA is extracted, prepared for poly(A)+ RNA sequencing, and FLARE^6^ is used to identify statistically significant edit clusters on both samples. **C)** The total number of FLARE edit clusters detected for each RBFOX2-rBE construct, and **D)** the total number of distinct RNA species edited by each fusion. **E)** The canonical RBFOX2 binding-site motif ((U)GCAUG) is recovered as the top *de novo,* HOMER-derived motif among an equally sampled number of edit clusters for each of the four RBFOX2-rBE chimeric proteins tested. **F)** For all RBFOX2-rBE fusions, the fraction of replicable clusters containing the canonical RBFOX2 binding site is significantly higher than the fraction for each of the thirty randomly selected edit cluster sets chosen from the respective mRNA target exons and intron sequences (“shuffled clusters”). The enrichment (right) is calculated as the actual value divided by the mean of the shuffled values. **G)** A density plot of the distance between the replicable peak centers produced by RBFOX2-rBE fusions and the closest canonical RBFOX2 binding-site motif (top) demonstrates far higher enrichment near peak centers than an equivalent plot for rBEs not fused to RBFOX2 (enzymes alone, bottom). **H)** Across RBFOX2-rBE fusions, the fraction of replicable edit clusters overlapping confident RBFOX2-APOBEC1 eCLIP peaks are significantly higher than the fraction for each of the thirty randomly selected peak sets chosen from the respective mRNA target exons and intron sequences (“shuffled clusters”). As in 2F, enrichment is calculated as the actual value divided by the mean of the shuffled values. **I-J)** Relationship between sets of I) edit clusters and J) RNA species detected by each RBFOX2- rBE fusion. A grey dot represents each RBFOX2-rBE set for each relationship considered. The number of I) edit cluster or J) RNA species values in each union is represented by a black bar. Vertical lines bisect colored dots to connect RBFOX2-rBEs with intersecting values, and single-colored dots without a line indicate unique clusters for the respective RBFOX2-rBE. **K)** The canonical RBFOX2 binding-site motif ((U)GCAUG) is recovered as the top *de novo,* HOMER-derived motif among edit clusters uniquely recovered by RBFOX2-rBE fusion.

Editing-based detection provides a larger temporal window of interaction capture (72 hours in our experiments), whereas eCLIP offers a momentary snapshot of RBFOX2 interactions. Also, we expect edits up to 200 base pairs away from eCLIP-based RBP-RNA interaction sites^41^ (for a more in depth analysis, please see refs.^6,41^). Notably, RBFOX2-APOBEC1 displays a higher frequency of capturing the RBFOX2 motif with the upstream U present in some RBFOX2 targets ((U)GCAUG) compared to other RBFOX2-rBE fusions (as depicted in Figure 2E). However, it is important to consider that APOBEC1 prefers editing substrate bases surrounded by A/U-rich regions^7^. This preference may explain the more frequent occurrence of the upstream U in the motifs enriched by RBFOX2-APOBEC1.

Subsequently, we assessed the degree of overlap among RBFOX2-rBE edit clusters originating from various rBEs. Intriguingly, only 99 clusters exhibited commonality across all fusions (Figure 2I and J). Even though the clusters detected by all fusions displayed a notable enrichment for the ((U)GCAUG motif (as shown in Figure 2K), it is noteworthy that the clusters unique to -8e, -A2dd (R), and -APOBEC1 RBFOX2 fusions also exhibited statistically significant enrichment for the (U)GCAUG motifs. This suggests that each enzyme introduces edits at distinct RBFOX2 binding sites (as illustrated in Figure 2K and Supplementary Figure 2D). Given that the RBFOX2 fusions solely differ by the rBE, our findings imply that enzyme-specific editing biases can lead to varying editing frequencies at distinct segments of the same RNA species bound by RBFOX2.

### Sequence context preferences of rBEs influence detection of sequence-specific RBP binding sites

To gain deeper insights into the intrinsic sequence preferences of the rBEs and how their fusion with sequence-specific RBPs influences their editing patterns at binding sites, we conducted a detailed analysis of the sequences surrounding the edits within the clusters derived from the RBFOX2-rBE fusion experiments. We employed a rigorous approach, training eight distinct two-layer convolutional neural networks (CNNs) to delve into the unique characteristics of each of the four rBEs individually, as well as their behavior when fused with RBFOX2. For each model, we used one-hot encoded 200-bp sequences from clusters associated with a specific rBE or RBFOX2-rBE fusions as positive examples, while an equal number of one-hot encoded 200- bp sequences from all other rBEs or RBFOX2-rBE fusions, respectively, served as negative examples. Each model was trained to provide a binary prediction, determining whether a given region was edited using that specific enzyme. To ensure effective learning without overfitting, we continued training until there was a minimal marginal reduction in the binary cross-entropy loss score on a separate validation dataset.

Across 8e, A2dd (R), APOBEC1, and 7.10 (V82G), these models can distinguish clusters generated by their respective target rBE from those produced by other rBEs fairly well, with the rBE-only variations on the whole performing better fusions (AUCs of 0.715, 0.839, 0.787, and 0.669, respectively) than those for the rBE fusions (AUCs of 0.701, 0.696, and 0.721) (Figure 3A). Interestingly, models trained on clusters generated solely by rBEs exhibited similar performance when evaluated on RBP-rBE fusion clusters, compared to models trained directly on RBP-rBE fusion clusters (Figure 3B). These observations suggest that while fusion with an RBP can alter the editing profile of an rBE, the underlying rBE-specific biases can persist. Consequently, even when different RBFOX2-rBE fusions associate with the same binding sites, the composition of the surrounding sequence at the RBP binding sites, along with the inherent rBE preferences, jointly influence whether the binding event will result in edits.

**Figure 3:**
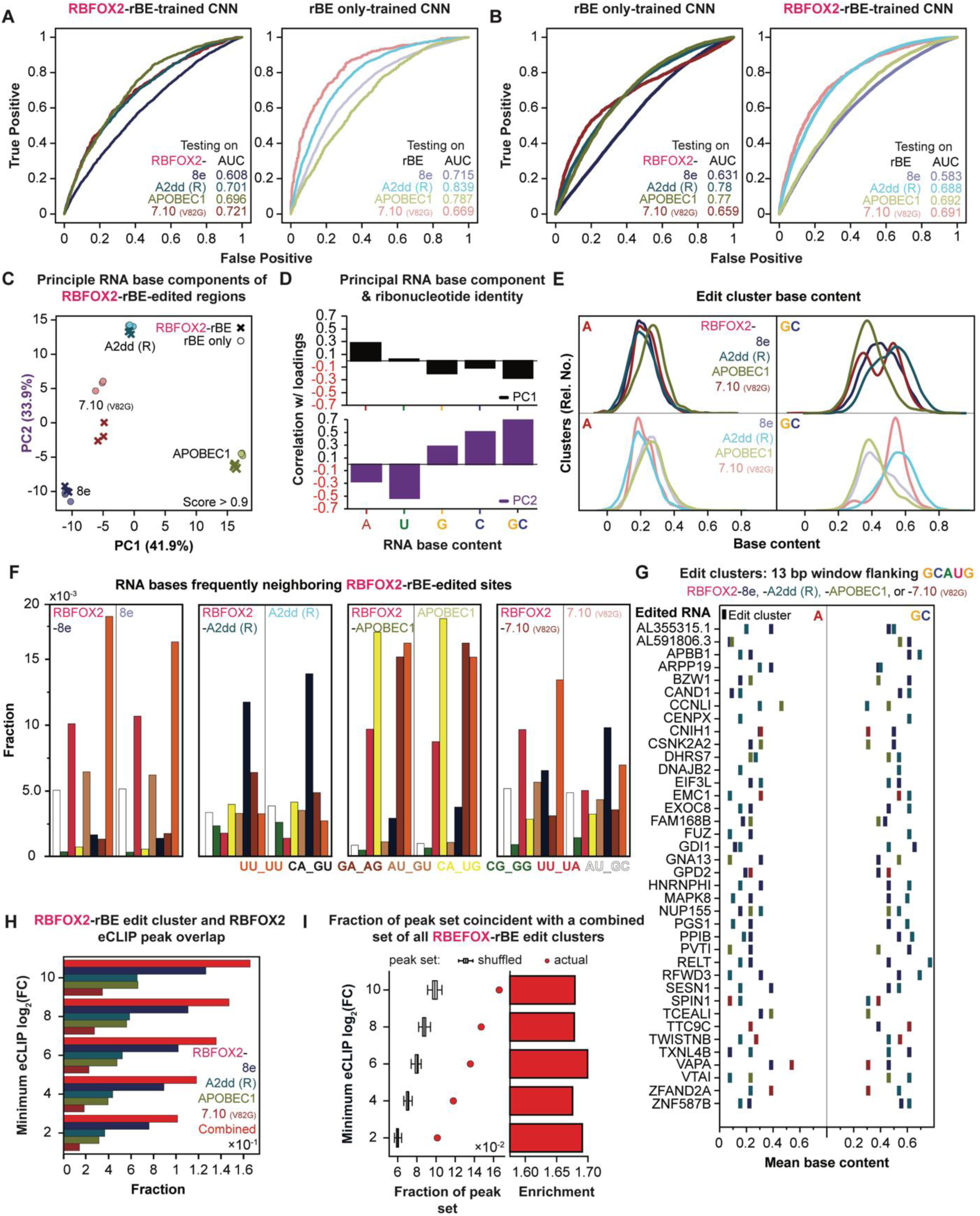
RBFOX2-rBE and rBE only bias. **A)** A simple (two convolutional layer) CNN discerns enzyme-specific bias in clusters from the enzyme-RBFOX2 fusion experiments and the enzyme alone experiments; AUCs for the enzymes alone are generally higher, implying more readily identifiable bias. **B)** Left, a CNN trained to recognize bias in the enzyme-alone clusters predicts fusion peak enzyme identity well and, in three out of four cases, better than the CNN trained on the fusion clusters themselves. Right, a CNN trained on enzyme-fusion clusters performs well (based on AUC) on enzyme-only clusters. However, they are not as well as the training-testing combinations depicted in the chart on the left. **C)** Edited sites have distinct and highly replicable enzyme-specific flanking base context preferences – whether alone or fused to RBFOX2 – as demonstrated by clear separation on a PCA plot of flanking context counts. **D)** The first and second principal components from the PCA plot in C exhibit strong and distinct contributions from each RNA base (A, U, G, or C) or combination (GC). **E)** A density plot of the overall peak adenosine (A, left) and guanosine-cytosine (GC) content for RBFOX2-rBE (top) or the fusion without RBFOX2 (rBE only, bottom), which are consistent between enzyme alone and enzyme fusion for any given enzyme. **F)** Edit site context preferences diverge in an enzyme-specific manner consistent between rBE- only and RBFOX2-rBE edits. **G)** On genes with two GCAUG core motif-containing clusters edited by two different enzymes, the combination of peak base contents and enzyme editing context specificities depicted in panel E dictate which region is more likely to be edited by each enzyme. **H)** The fraction of combined edit clusters (red bar) and individual edit cluster sets (see key) that intersect with RBFOX2-APOBEC1 eCLIP peak sequences (sites). **I)** The fraction of the RBFOX2-APOBEC1 eCLIP peak sequences for peaks above a given minimum log2(fold change) (Y-axis) that overlap with sequences of a combined set of all replicable RBFOX2-rBE edit clusters. Boxplots denote the overlap between each of thirty randomly selected artificial eCLIP peak sets and the combined edit cluster set, while the red circles show the overlap of the actual RBFOX2-APOBEC1 eCLIP peaks. The enrichment (right) is calculated as the actual value divided by the mean of the shuffled values.

We next aimed to uncover which specific sequence features underlie these differences between the preferred editing contexts of each rBE. To do so, we characterized the four bases flanking high confidence edit sites (SAILOR score > 0.9) for each rBE. A principal components analysis of the counts of each flanking base context within each RBP-rBE fusion dataset – based on proximity in PCA space along the first two principal components (PCs) – indicates that each rBE has highly replicable distributions of flanking base pairs, and that edits generated by the same rBEs (whether alone or fused to RBPs) are found in more similar contexts to one another than to those generated by any other rBE or RBP-rBE fusion (Figure 3C). These results support the conclusions from the CNN-based analyses described above (Figure 3A, B). We examined the loadings for the contribution from each possible flanking base context with respect to PCs 1 and 2, and found that the factor loadings for PC1 exhibited strong positive contribution from A/U bases and negative contribution from G/C bases (Figure 3D). The opposite was true for PC2, as it had negative contribution from A/U bases and positive contribution from G/C (Figure 3D). Indeed, APOBEC1 and RBFOX2-APOBEC1 flanking contexts have high PC1 (A/U) values and low PC2 (G/C), consistent with the known APOBEC1 preference for cytosines flanked by As and Us^7,42,43^ and its low tolerance for Gs^8^, respectively. These rBE-specific affinities are evident at the cluster-wide level, as shown by similarly varying cluster-level base contents (Figure 3E, and Supplementary Figure 3A). We see that 7.10 (V82G) and A2dd (R) clusters tend to have higher GC content than APOBEC1 and 8e clusters, and APOBEC1 clusters are enriched for A content compared to clusters from other enzymes. Further examination of the distribution of edit flanking contexts revealed that out of the 16 possible flanking bases, 12 were either A’s or U’s. This reflects A/U flanking context bias, consistent with APOBEC1’s preferences^7,42,43^. In contrast, A2dd (R) edit sites exhibited an enrichment for flanking contexts involving G’s and C’s, indicating a higher tolerance for these bases (Figure 3F).

The implications of these varying preferred contexts for different RBP-rBE fusions are that authentic RBP binding sites, depending on their specific nucleotide contexts, may be better suited for editing by one RBP-rBE fusion over another. In other words, no single RBP-rBE fusion can edit the entire universe of existing binding sites. Pairs of core RBFOX2 binding sites (GCAUG) on the same gene are sometimes preferentially edited by different rBEs (Figure 3G), with the nucleotide context of the 4 bases flanking the core GCAUG motif serving as a predictive element for which rBE is most likely to edit at each site. For example, when we focus on the FLARE- derived significant edit clusters found on CCNLI RNA, we notice that the RBFOX2 binding site with a greater GC content is only significantly edited by RBFOX2-A2dd (R), and the site with higher A content is only significantly edited by the RBFOX2-APOBEC1 fusion (as shown in Figure 3G and Supplementary Figure 3B).

These rBE-specific biases imply that, at least for RBFOX2, a single RBP-rBE fusion is insufficient to capture the entire spectrum of RBP-RNA interactions. Combining multiple rBEs is expected to provide better coverage of true RBP binding sites than any individual fusion. To test this, we merged the confident cluster sets from all RBFOX2-rBEs into a single combined set and assessed the fraction of overlap with RBFOX2-APOBEC1 eCLIP peaks (Figure 2C and 3H). Through empirical permutation testing, we observed that the combined cluster set captured a greater fraction of RBFOX2 eCLIP peaks than any of the individual fusions, and this overlap was statistically higher than expected by random chance (Figure 3H, I). Importantly, even though 8e had higher editing activity, less active enzymes still captured eCLIP clusters that 8e missed (Figure 3H), indicating that increasing the editing efficacy of certain rBEs may yield more clusters, but there will always be some subset of binding targets that remain undiscoverable without the use of other enzymes with editing context preferences better suited for those targets.

Lastly, we assessed whether the editors exhibited editing bias towards specific RNA regions. We found that, while most rBEs, both as fusions to RBFOX2 and individually, generally favored editing in 3’UTRs, 8e and A2dd (R) exhibited slightly more editing in coding regions (CDS) (Supplementary Figure 3C). In conclusion, the suitability of rBEs to fuse with sequence-specific RBPs is influenced by sequence context, and a combination of multiple rBEs is recommended for achieving higher coverage and discovery of RBP-RNA binding sites.

### Ribosome-subunit RPS2-rBE fusions robustly detect transcriptome-wide mRNA translation changes

We previously demonstrated that fusion of APOBEC1 to the core small ribosomal subunit protein RPS2 (RPS2-APOBEC1 or RiboSTAMP) enabled the measurement of ribosome-mRNA interactions, even in single cells^5^. Here, we evaluated RPS2 fusions to 8e, A2dd (R) and 7.10 (V82G), in comparison to APOBEC1 (Figures 4A and 4B). Plasmids encoding RPS2-rBE fusions were transfected into HEK293XT cells and induced with doxycycline for 24 hours, after which cells were treated with either dimethyl sulfoxide (DMSO) vehicle (-) or 100 nM of the mTOR pathway inhibitor Torin-1 (+) for 48 hrs (Figure 4B) to induce changes in mRNA translation. The number of edits per sequencing read (edits per read or “EPR”) for each experiment was measured as a proxy for mRNA translation. We conducted three experiments for each RPS2-rBE fusion and concentrated our analyses on mRNAs that were consistently edited in all biological replicates. As expected, we observed generally lower EPR values in cells treated with Torin-1 compared to DMSO treatment for all RPS2-rBEs (Figure 4C). We also determined that more mRNAs exhibited statistically significant decreases in EPR than increases (Figure 4D). Notably, the Torin-1- mediated reduction of editing by the RPS2-rBE fusions was more evident among 5’ terminal oligopyrimidine tract (TOP)-containing mRNAs^44^ (Figure 4E). Among the tested enzymes, the RPS2-A2dd (R) showed the lowest p-value and the largest t-test statistic for decreasing EPR in (TOP)-containing mRNAs between Torin-1 and DMSO conditions (t-test statistic 14.84, p-value<10^-44^ ; Figure 4F), followed by 8e (t-test statistic 9.43, p-value <10^-19^), APOBEC1 (t-test statistic 4.24, p-value<10^-4^) and 7.10 (V82G) (t-test 3.44, p-value<10^-3^) (Figure 4E, 4F). Notably, while all RPS2-rBE fusions similarly detected the majority of the fifty TOP-containing mRNAs, the overlap for non-TOP-containing RNAs that experienced significantly reduced editing exhibited a smaller overlap (Supplementary Figures 4A, B).

**Figure 4:**
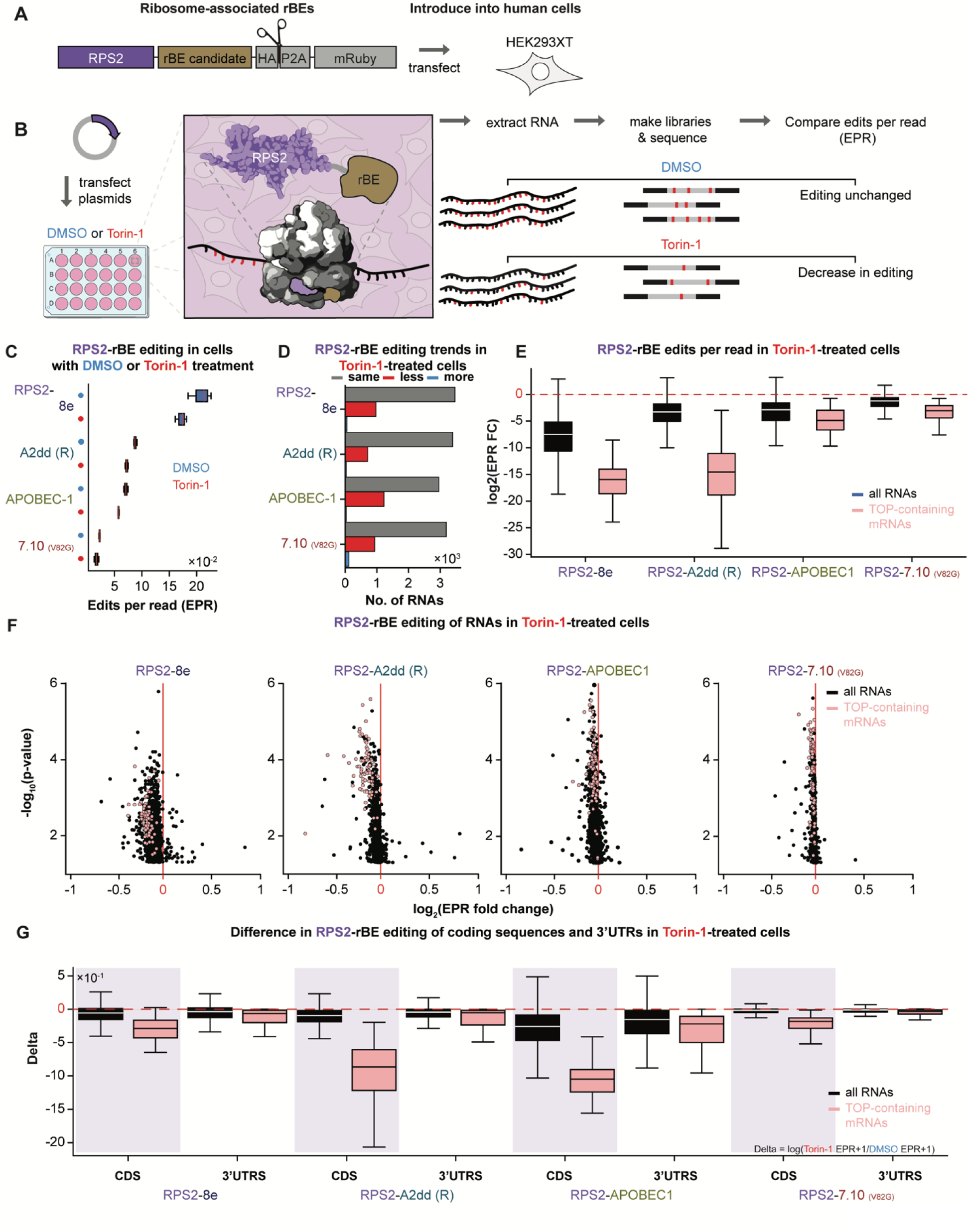
RNA editing as a proxy for translation with RPS2, the top rBE candidates. **A)** Construct bearing the small ribosomal subunit protein, RPS2 (purple) fused to a candidate rBE (brown) at its C-terminal end (“Translation-directed editor”). **B)** Strategy to detect RPS2-directed editing. A plasmid encoding the construct in A is transfected into HEK293XT cells, and cells are incubated with Torin-1 to inhibit translation or with a DMSO vehicle as control 48 hrs. The RPS2-rBE fusions are predicted to incorporate into the ribosome for translation-meditated mRNA editing. After the treatment, libraries are prepared from isolated total poly(A)+ RNA and sequenced. Torin-1 treatment is predicted to cause a reduction in editing, as judged by the edits per read (EPR) in the sequencing reads. **C)** Overall RPS2-rBE edits per read (EPR) in cells treated with Torin-1 are reduced compared to EPR in untreated (DMSO) cells across all RPS2-rBE fusions. **D)** Number of RNAs experiencing statistically significant decrease (red) or increase (blue) in RPS2-rBE EPR in Torin-1-treated cells vs. DMSO-treated control, as well as RNAs that exhibited no significant change (grey). **E)** The log2-transformed Torin1-mediated decrease in EPR, calculated as EPR in the torin1 condition divided by the EPR in the untreated state, compared across all RPS2-enzyme fusions for poly(A)+ RNAs (black) and TOP-containing mRNAs (pink). **F)** Edits per read difference (fold-change, x-axis) between Torin-1- and DMSO-treated cells and the associated statistical significance (log2-transformed *p*-value, y-axis) for each RNA (dots). Values for Poly(A)+ RNAs (black) and TOP-containing mRNAs (pink) are shown. **G)** The difference between edits per read between Torin-1- and DMSO-treated cells when EPR analyses are focused on either coding sequences (CDS) or 3’ untranslated regions (UTRs) indicate changes in editing are specific to the CDSs for both poly(A)+ RNAs (black) and TOP-containing mRNAs (pink). Here, as with 4E-F, differences are more conspicuous among TOP- containing mRNAs.

We previously observed RPS2-APOBEC1 editing within 3’UTRs^5^. We compared the relative editing rates between mRNA coding sequences (CDSs) and 3’UTRs across our experimental conditions for our rBE-candidates. When considering all mRNAs, the ratios of EPR in CDS regions to 3’UTRs were similar across enzymes for mRNAs from both untreated and Torin-1 treatment conditions (Figure 4G, compare the black CDS boxplot to the black 3’UTR boxplot for each individual enzyme). However, in TOP-containing genes, the CDS to 3’UTR editing ratio skews higher for each individual enzyme, indicating stronger decrease in editing among CDS regions compared to 3’UTRs (Figure 4G, compare pink CDS boxplot to pink 3’UTRs boxplot for each individual enzyme). The ratios of the means of CDS/3’UTR editing as depicted in the boxplot are as follows: RPS2-8e = 1.873, RPS2-APOBEC1 = 1.384, RPS2-7.10 (V82G) = 2.331, and RPS2-A2dd (R) = 2.456. Since TOP genes are among the most highly translated genes, the increased propensity for CDS edits may indicate higher ribosome load on the coding sequences of these transcripts in standard growth conditions. This interpretation is bolstered by the observation that the CDS to 3’UTR editing ratio of TOP genes is lower in Torin-1 treated cells (Figure 4G, and Supplementary Figure 4D). Therefore, as with RPS2-APOBEC1, other RPS2- rBE fusions demonstrate editing in 3’UTRs, with RPS2-mediated editing more pronounced in CDS regions for all rBEs.

Like our observations for RBFOX2-rBE fusions, our analysis of the flanking base context for edits revealed distinct base context preferences for each enzyme fusion (Supplementary Figure 4E, F) which are unaltered by the effects of Torin-1, as Torin-1 replicates clustered together with untreated replicates (Supplementary Figure 4E). We observed that the loadings for PC1 had strong positive contribution from A/U bases, and strong negative contribution from G/C bases (Figure 4E, G). Conversely, the PC2 loadings had strong negative contributions from A/U and strong positive contributions from G/C bases (Supplementary Figure 4E, G). These influences are also reflected in the bases that often neighbor edited sites (Supplementary Figure 4G). However, as ribosomes are recruited to a substantially broader sequence area than sequence-specific RBPs like RBFOX2, we find that single rBE fusions to ribosome components is sufficient to measure general aspects of mRNA translation. To detect variations in RPS2-mediated editing rates, we suggest using 8e, A2dd (R), or APOBEC1. This is because the fold decreases in editing are more pronounced in Torin-1-treated cells compared to DMSO-treated cells with these RPS2- rBE fusions as opposed to the less sensitive RPS2-7.10 (V82G) fusion (Figure 4F,G).

### Evaluating combinations of C-to-U and A-to-I rBEs to assay dual editing compatibility

A-to-I and C-to-U edits can be used simultaneously to interrogate the binding of two distinct RNA-binding proteins on the same RNA transcript (as with TRIBE-STAMP^45^ that uses hAcd (hyperTRIBE) and APOBEC1). To test this possibility for our rBE fusions, we modified the synthetic 3’ UTR in our twelve MS2 stem-loop reporter to also include PP7 stem-loops (Figure 5A, left). The PP7 bacteriophage coat protein (PP7-CP) binds to the PP7 stem-loops, which coupled with the MS2 stem-loops can simultaneously recruit both C-to-U and A-to-I editing enzymes to the same 3’UTR. Different distributions of the MCP and PP7-CP binding sites on the reporter can also yield insights into the effects of binding site proximity on RNA co-editing (Figure 5A, left). To create PP7-CP fusions, we replaced MCP with PP7-CP in the MCP-APOBEC1 plasmid and substituted APOBEC1 with either precise (7.10 (V82G)) or more active (8e) A-to-I editors selected based on enrichment scores and noise levels from RBFOX2 fusion and reporter experiments. After, plasmids encoding the MCP-APOBEC1, one of the PP7-CP-A-to-I fusions (PP7-CP-7.10 (V82G) or PP7-CP-8e), and one of the reporter mRNAs were co-transfected into HEK293XT cells (Figure 5A, B) in biological duplicates. The editing experiments were then carried out as for our initial rBE screen (Figure 5A, B).

**Figure 5:**
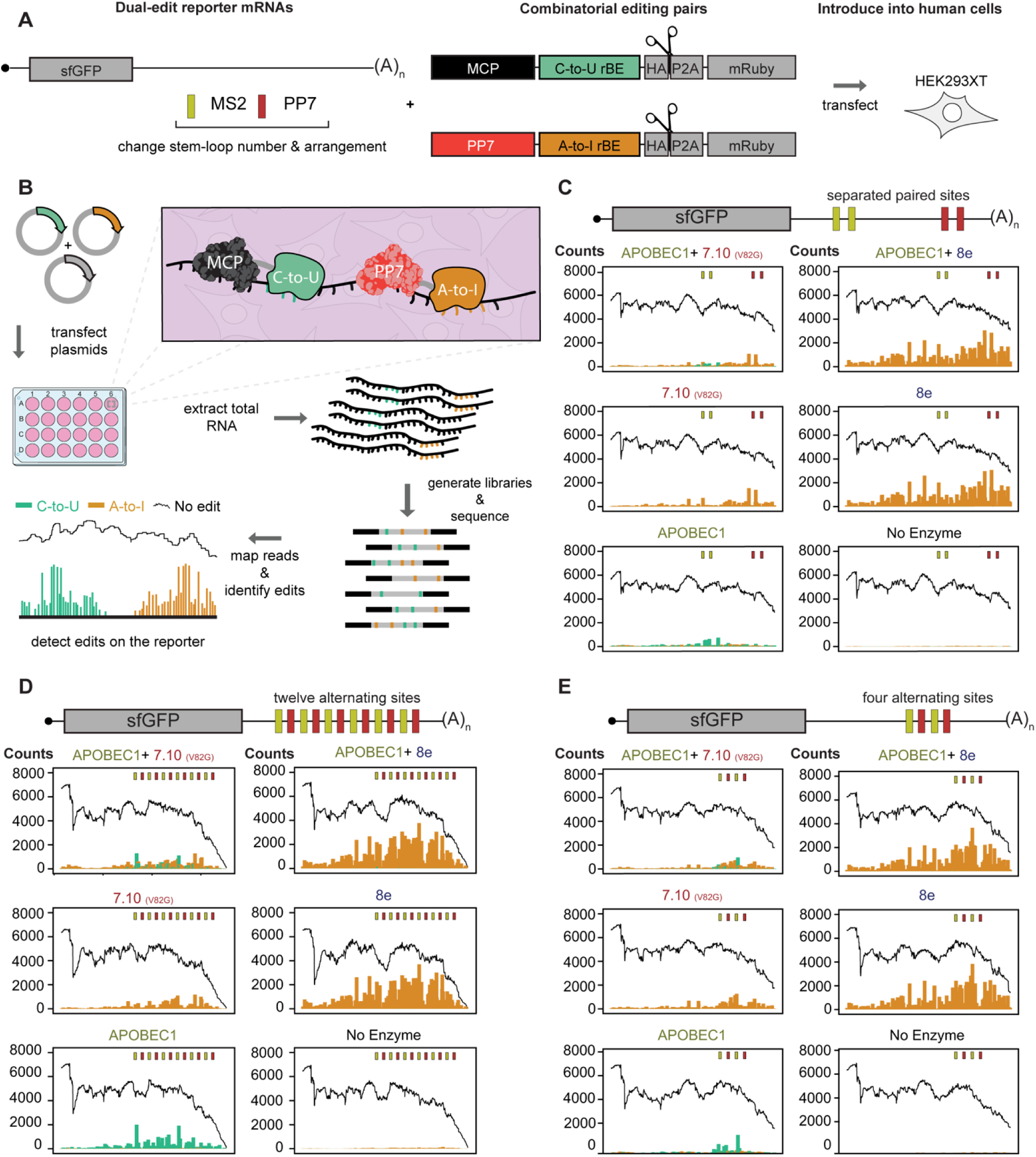
A combinatorial editing reporter system to identify multiple RBP associations with the same transcript. **A)** Components that are used to test combinatorial C-to-U and A-to-I rBE pairs. The system uses a reporter mRNA with varied MS2- and PP7- stem-loop distributions in the 3’ UTR. The reporter binds the MS2 and PP7 coat proteins fused to C-to-U and A-to-I rBEs. **B)** A combinatorial editing strategy. Plasmids encoding C-to-U and A-to-I fused MCP and PP7 proteins are co-transfected into HEK293XT cells with the reporter bearing both binding sites. After, the edits are detected on the reporter with targeted RNA sequencing. **C-E)** Distribution of C-to-U (green) and A-to-I (orange) edits deposited by each of five different enzyme combinations and the reporter without enzymes. The edits are mapped along each of the three distinct reporters. One reporter bears a **C)** MS2 (yellow) and PP7 (red) stem-loops space 50 bp apart, with 350 bp separating the pairs. The other two reporters contain **D)** twelve or **E)** four alternating MS2 and PP7 binding sites spaced 50 bp apart.

Our dual-editing reporter system revealed distinct co-editing patterns. MCP-APOBEC1 and PP7-CP-7.10 (V82G) deposited edits near their respective binding sites in all the MS2 and PP7 stem-loop distributions tested but are more conspicuous in the constructs in which we alternated the binding sites (Figure 5C-E, and Supplementary Figure 5C-E). Moreover, A-to-I editing was dominant when MCP-APOBEC1 was paired with PP7-CP-8e (Figure 5C-E, and Supplementary Figure 5C-E). Indeed, although the MCP-APOBEC1 could deposit detectable C-to-U editing when the A-to-I editor was omitted, co-expressing the enzyme with PP7-CP-8e led to a marked reduction of C-to-U edits (Figure 5C-E, and Supplementary Figure 5C-E). Further, PP7- CP-8e A-to-I editing looked similar whether the fusion was co-expressed with MCP-APOBEC1 or not (Figure 5C-E, and Supplementary Figure 5C-E). Lastly, the PP7-CP-8e also yielded a higher rate of edit spillover into the neighboring MS2 binding sites than did MCP-APOBEC1 and PP7- CP-7.10 (V82G) (Figure 5C-E, and Supplementary Figure 5C-E).

MCP-APOBEC1 and PP7-CP-7.10 (V82G) deposited fewer edits when co-expressed than when expressed on their own, consistent with our observation where one editor (8e) impacted the editing of the other (APOBEC1, Figure 5C-E, and Supplementary Figure 5C-E). Further, as with our 12X MS2-SL reporter, MCP-APOBEC1 and PP7-CP-7.10 (V82G) produced less edit spillover than PP7-CP-8e (Figure 5C-E, and Supplementary Figure 5C-E). Our reporter demonstrates the precision of the 7.10 (V82G) and APOBEC1 pair with paired MS2 and PP7 binding sites that were 350 bases apart. On this reporter, MCP-APOBEC1 and PP7-CP-7.10 (V82G) edits clustered near their corresponding binding sites, while PP7-CP-8e editing clusters near its binding site but produces much more evenly distributed edits across the 3’UTR (Figure 5C, and Supplementary Figure 5C). Further, reducing the number of twelve alternating MS2 and PP7 binding sites to four produced tighter editing patterns for MCP-APOBEC1 and PP7-7.10 (V82G) (Figure 5D, E, and Supplementary Figure 5D, E), highlighting the editing precision of this pair. We conclude and recommend that APOBEC1 and 7.10 (V82G) are a robust, balanced pair for dual-RBP-based editing applications.

## Discussion

We have introduced PRINTER, a systematic workflow of experimental and computational assays designed to comprehensively assess the capabilities of over thirty candidate RNA base editors (rBEs) for probing protein-RNA interactomes within living cells. Our RNA “tethering” assay effectively recruits rBEs to a synthetic 3’ UTR on a reporter mRNA, where they catalyze edits. Subsequently, we rigorously analyze on- and off-target editing activities using targeted and transcriptome-wide RNA sequencing. Our findings reveal a rich diversity in the sensitivity and specificity of the rBEs.

Among the candidates evaluated, we have identified seven promising rBEs that hold significant potential to expand the scope of RBP-directed editing. These noteworthy editors include 8e, evoAPOBEC1, A2dd (R), 7.10 (V82G), 7.10, an APOBEC3A mutant (A3A (Y132G/K30R)), and A2dd (R-S), as listed in Table 1. Notably, several of these editors exhibited improved signals when compared to TRIBE and hyperTRIBE enzymes, underscoring their enhanced performance in RBP interactome studies. Additionally, we have identified editors with varying editing activity levels, such as A2dd (R-S) with reduced activity and evoAPOBEC1 with enhanced activity compared to the APOBEC1 enzyme used in STAMP, thus broadening the spectrum of editing control for RBP-mediated interactions.

However, it is worth noting that some enzymes in our screening did not perform robustly in our reporter assay. Cas9-mediated DNA editors (in the absence of Cas9) like evoCDA1, evoFERNY, evoA3A, AID, APOBEC3G, and several AID variants were unable to edit RNA effectively. Additionally, SECURE rAPOBEC1 (R33A) and SECURE rAPOBEC1 (R33A, K34A), which had shown detectable RNA edits in previous studies, did not produce noticeable editing in our reporter RNA. Furthermore, ADARcd^4^ and hyperADARcd^15^ (referred to as hAcd in this study), the enzymes employed in TRIBE and hyper-TRIBE, respectively, failed to generate detectable edits in our experiments. This discrepancy in ADAR editing could be attributed to differences in enzyme sources, as the TRIBE enzymes originate from fruit flies, whereas the A2dd enzymes are human-derived. Another possibility is that dsRNA, which is the preferred substrate of ADAR^4,9–14^, is less abundant in the cytoplasm of cells, limiting the activity of these enzymes^46^.

Our RNA base editor tethering approach offers a rapid means to characterize enzymes before their application in RBP-mediated editing. It also enables the identification of DNA editors with reduced off-target RNA editing, addressing a critical aspect of the field’s current limitations. The challenge of achieving an optimal signal-to-noise ratio in RBP-directed editing experiments remains significant. Striking the right balance depends on the specific RBP-related problem at hand. In our studies, higher enzyme activity tends to result in more edits away from the binding sites, as exemplified by 8e in our reporter assays. While this may introduce some noise, it’s important to note that 8e captures a largely distinct set of genuine RBFOX2 targets. On the other end of the spectrum, utilizing enzymes that rely on infrequent sequences or structures, such as the dsRNA editing ADARs, may fail to encompass the full spectrum of protein-RNA interactions in conditions where these structures are rare, like the cytoplasm.

Our combined computational and experimental strategies represent a step toward addressing the limitations inherent in editing-based detection approaches. These strategies provide valuable insights into the inherent biases of rBEs and their impact on the detection of protein-RNA interactions. Specifically, the observed preferences of certain rBEs for distinct sequence contexts, such as the GC-rich sequences preferred by A2dd (R), 7.10 (V82G), and 8e, or the A-rich contexts preferred by APOBEC1, offer valuable guidance for selecting the most suitable enzyme for a given study. Additionally, considering factors like 3’UTR bias and precision of editing helps researchers make informed decisions when choosing an rBE for their experiments. Specifically, 8e and A2dd (R) appeared to have less 3’UTR bias, which may make them more suitable for studies where RBPs associate with regions like the CDS. Precision was highest among 7.10 (V82G), 8e, and then A2dd (R) in those studies. We recommend using 7.10 (V82G) where GC tolerance and precision are required and 8e or A2dd (R) to maximize the number of recovered RNA targets and edit clusters.

Our study emphasizes the importance of acknowledging and addressing enzyme bias when designing experiments and interpreting their outcomes. Relying solely on one enzyme can lead to false negatives and an incomplete understanding of protein-RNA interactions. A case in point is our RBFOX2-rBE fusions, which produce edit clusters that overlap with genuine yet largely distinct sets of RBFOX2-APOBEC1 eCLIP targets. Thus, our recommendation to assay RBPs with multiple enzymes ensures a more comprehensive capture of genuine targets. These datasets can subsequently be analyzed to identify common sequence motifs that persist across different enzymes, facilitating the discovery of conserved binding sites. Furthermore, our work challenges the assumption that a single enzyme can universally address any type of protein-RNA interaction. The varying performance of different enzymes in different contexts underscores the need for a diversified toolbox of rBEs. Enzymes should be chosen based on their compatibility with the specific experimental goals, whether that involves GC-rich sequences, precision editing, or the detection of distinct types of interactions. To illustrate this point, consider RBFOX2-A2dd (R), which results in only a modest enrichment of RBFOX2 motif sequences and RBFOX2 eCLIP peaks. In contrast, RPS2-A2dd (R) outperformed all others in detecting Torin-1-mediated translational repression. This scenario underscores the ongoing need for the field to expand the repertoire of available enzymes for RBP-directed editing, a pursuit exemplified by the recent introduction of TadA-CD C-to-U editors, which are derived from the TadA-8e (8e) A-to-I editor profiled in our study ^25^.

Our findings also highlight the potential for combining multiple rBEs to achieve more comprehensive results or evaluate which RBPs are bound to the same RNA molecule. We find that only some enzymes are compatible with “dual editing” on the same RNA^45,47,48^. Compatibility is important for the interpretability of the results because mismatched enzyme activities could lead to one enzyme masking the edits of the other. We recommend APOBEC1 (C-to-U) and 7.10 (V82G) (A-to-I) which captures co-binding across three distinct RBP binding site configurations. As with our single RBP studies, using more than one C-to-U and A-to-I enzyme pair would likely yield a more complete view of RBP co-binding than using a single pair alone. Thus, future work in identifying RNA base editors with different activities will also enable the field to identify more useful co-editing pairs.

In summary, our study not only addresses key limitations in the field of RNA base editor-mediated detection of protein-RNA interactions but also presents opportunities for further refinement and expansion. By modifying the sequences in the linker regions of our reporter (currently 50bp RNA linker flanking MS2 stem-loops), we can evaluate enzymes with distinct sequence context preferences, paving the way for engineered rBEs, as has been previously conducted with DNA editors^8,26,27^. Additionally, there is vast potential to evaluate the ability of rBEs to detect localized RNAs^49,50^. For example, integrating the molecular recording strategy, localized RNA recording, and proximity-specific ribosome profiling data from yeast added insights into the interplay between RNA localization and translation at the ER and mitochondria that is impossible with either method alone^49–52^. Adapting optimized rBEs to identify localized RNAs can expand on the localized RNA recording strategy and complement methods like APEX-Seq and proximity-specific ribosome profiling^49–53^. Here, distinct editors may also be used to record RNA molecules that interact with multiple sub-cellular locales, as previously suggested with combined localized RNA recording and TRIBE^49,50^.

## Materials and Methods

### Cloning

#### 12x-MS2 stem-loop mRNA reporter

The pcDNA3.1 (-) Mammalian Expression Vector (Invitrogen, Cat # V79520) was digested with the NheI (Thermo Fisher Scientific, Cat # FD0974) and the MssI (PmeI, Thermo Fisher Scientific, Cat # ER1341) restriction enzymes. After, fragments (IDT gene blocks) bearing the human codon-optimized (IDT tool) super folder green fluorescent protein^54^ (sfGFP) coding sequence and a synthetic 3’ UTR made of twelve version-six MS2 bacteriophage stem-loops^8^ (12XMBSV6) were cloned into the digested pcDNA3.1 (-) vector (Thermo Fisher Scientific, Cat # V79520) via Gibson assembly^55^.

### MCP-RNA Base-Editor (rBE) Fusions

#### Strategy 1: Gateway cloning

The pDONR221 Gateway vector (Thermo Fisher Scientific, Cat # 12536017) was first digested with AfiII (New England Biolabs, Cat # R0520S) and EcoRV (New England Biolabs, Cat # R0195S) restriction enzymes to remove the attP1, ccdB, cmR, and attP2 cassettes. After, two gene fragments bearing the MS2 coat protein-coding sequence flanked by the attL1 and attL2 sequence were cloned into the digested vector via Gibson assembly. The resulting MCP- pDONR221, pHCMM14, was then used to insert the MCP into each rBE-bearing destinations vector using Gateway LR cloning^56^.

### Preparation of destination vector for Gateway LR cloning

The pLIX403_Capture1_APOBEC_HA_P2A_mRuby (Plasmid #183901) vector^5^. was digested with PspXI (New England Biolabs, Cat # R0656S) and BstZ17I-HF (New England Biolabs, Cat # R3594S) to remove the APOBEC1 coding sequence. The digested fragments were then cleaned using the QIAquick PCR Purification Kit (Qiagen, Cat # 28104) and eluted in 18 μL water. Afterward, 2 μL of E-Gel Sample Loading Buffer, 1X (Thermo Fisher Scientific, cat # 10482055), were added to the eluate, and 20 μL of the mix was loaded onto a well on a 2% Agarose E-gel (Thermo Fisher Scientific, Cat # G401002). Next, the cassette was loaded onto the E-Gel Power Snap Electrophoresis Device (Thermo Fisher Scientific, Cat # G8100) and run for 13 minutes on the 1-2% agarose gel setting. Finally, a band corresponding to the size of the backbone vector without the APOBEC1 sequence was excised from the agarose gel and purified using the Qiagen mini-elute gel extraction kit (Qiagen, Cat # 28604).

### PCR-amplification of rBE coding sequences

The open reading frames encoding distinct rBE candidates were amplified using Q5 High-Fidelity DNA Polymerase (New England Biolabs, Cat # M0491). The PCR primers were designed to produce PCR product flanked by ∼ 40 bp of sequence complementary to the target backbone vector, a requirement for Gibson assembly. The PCR products were then purified and quantified for the digested backbone vector (as described above for digested vector purification).

### Gibson assembly of PCR-amplified rBE CDSs with the STAMP backbone vector

Gibson assembly reactions were assembled by adding 200-400 ng of each purified amplicon and ∼50 ng of digested backbone vector together with 15 μL of 2X Gibson master mix and molecular grade water to a total of 20 μL volume. The reactions were then incubated at 50

°C for 50 minutes. After, 2 μL of the reaction were transformed into MultiShot™ FlexPlate TOP10 Competent Cells (Thermo Fisher Scientific, Cat # C4081201). Successful cloning was confirmed by Sanger sequencing. Finally, the Gibson reactions produced destination vectors bearing distinct rBE candidates.

### Gateway cloning to generate MCP-rBE fusions

The entry clone plasmid bearing the MCP coding sequence (pHCMM14) was combined with each of the destination vectors carrying distinct rBE candidates (see above) using Gateway LR cloning (Thermo Fisher Scientific, Cat # 11791020). Next, 1 μL of the reaction was transformed into *E. coli* and the correct clones were confirmed by Sanger sequencing. The resulting plasmids were then transiently transfected into human embryonic kidney 293 cells (HEK293XT, see below).

### Strategy 2: Gibson assembly to generate MCP-rBE fusions

We switched to Gibson assembly to speed up the generation of MCP-rBE fusions, which would skip the gateway cloning step. First, the MCP-APOBEC1 fusion generated above with Gateway cloning was digested with the AfeI (New England Biolabs, Cat # R0652S) or FastDigest Eco47III (Thermo Fisher Scientific, Cat # FD0324) and PspXI (New England Biolabs, Cat # R0656S) restrictions enzymes to remove the APOBEC1 from downstream of the MCP. The coding sequence for each of the remaining enzymes in our panel was then amplified using primer that yielded fragments flanked by ∼40 bp sequences complementary to the digested backbone vector. The vector and PCR products were then purified as described above. After ∼50 ng of the digested vector was combined with 200-400 ng of the PCR-amplified rBE sequences, 15 μL of Gibson master mix, and water to bring the reactions to 20 μL each. The reactions were then incubated at 50 °C for 50 mins. After, the 2 μL of the Gibson reactions were transformed into E. coli and the clones were isolated from the resulting colonies and confirmed with Sanger sequencing. This approach yielded the remaining MCP-rBE fusions described in this report.

### Transient transfection of HEK293XT cells

Plasmids were transfected into HEK293XT cells using Lipofectamine 3000 (Thermo Fisher Scientific, Cat # L3000015). To prepare HEK cells for transfection, they were first grown in a 10 cm dish to 80% confluency. After, the media was removed, and the cells detached from the plate via TrypLE (Thermo Fisher Scientific, Cat # 12604039) treatment at 37 °C for ∼3 mins and subsequent pipette-mixing with 10 mL of DMEM (Thermo Fisher Scientific, Cat # 11965092) + 10% FBS (Thermo Fisher Scientific, Cat # 26140-079). The suspension was then transferred to a 15 mL conical tube, and cells were pelleted at 200 RCF for 5 minutes. The cells were resuspended in 3-5 mL of DMEM, and 20 μL of cell suspension was mixed with 20 μL of Trypan Blue (Thermo Fisher Scientific, Cat # 15250061). 10 μL of the mixture was then loaded onto a Dual-chamber Cell Counting Slide (Bio-Rad, Cat # 1450011), and the cells were counted using a Bio-RAD TC20 Automated Cell Counter. After cells were diluted to 174 cells/μL, adding 500 μL of cell suspension yielded ∼87,000 cells per well of a 24-well plate. The cells were incubated overnight, after which DNA-lipid complexes were generated using lipofectamine 3000 within 24 hrs of plating. For 12-well plates, 1,250 ng of DNA was used to create lipid-DNA complexes, while 625 ng was used for 24-well plates. Once created, the complexes were added drop-wise to the wells making sure to cover as much area of the well as possible with the drops. The plates were then placed in a 37 °C incubator overnight. The next day, 500 μL of fresh media containing 2ug/ml doxycycline (Dox) and 1µg/mL puromycin was added to the existing media in each well. The cells were then incubated at 37 °C for 48 or 72 hrs. After, the media was removed via aspiration, and the cells recovered by TryPLe (500 μL) treatment for 2 mins at 37 °C, followed by resuspension in 1 mL of DMEM + 10% FBS. The cells were then pelleted via centrifugation at 200 RCF for 5 mins at room temperature, the supernatant removed, and pellets stored at -80 °C until analysis. Alternatively, the aspirated media was replaced by 300-600 μL of TRIzol. The plates were either covered with foil seals and stored at -80 °C until RNA extraction or DNA-free total RNA was immediately isolated from the lysate using the Direct-zol RNA Miniprep Kit (Zymogen, Cat # R2052) using the RNA Purification protocol. The RNA was then quantified using the NanoDrop UV-Vis spectrophotometer.

### Targeted Reporter RNA Sequencing

Experiments were performed in duplicate with an APOBEC1 positive control and a “reporter alone” negative, and libraries were prepared with the same amount of starting RNA material for all samples for qualitative comparison. To do so, 500 ng of total RNA was subjected to reverse transcription using a Superscript IV reverse transcriptase kit (Thermo Fisher Scientific, cat # 18090010) in 20 µL volume. 1 μL of the resulting cDNA was then subjected to PCR amplification using Q5 High-Fidelity DNA Polymerase (New England Biolabs, Cat # M0491L) and primers specific to the reporter mRNA. The resulting amplicon was then purified as described by the “DNA fragment purification” section below. 1 ng of the purified fragment was then used to prepare sequencing libraries with the Nextera XT DNA Library Preparation Kit (Illumina, Cat # FC-131-2002 or Cat # FC-131-1024; Indexes: Nextera XT DNA Library Preparation Kit (24 samples), Cat # FC-131-1024 or IDT for Illumina DNA/RNA UD, Cat # 20027213). The resulting libraries were then purified and normalized with either the Nextera XT kit or standard normalization. Equimolar amounts of each library were then pooled to make a 4 nM library. After, a 6.5 pM library was denatured and loaded onto a MiSeq reagent cartridge (Illumina, Cat # MS- 102-3003, MS-102-3001, MS-103-1002, or MS-103-1001) and sequenced on the Illumina MiSeq with either single-end 150 or paired-end 150 read format.

### DNA fragment purification

PCR amplification (two 50 μL reactions) and plasmid digest (three 50 μL) reactions were combined into a single Eppendorf tube. The reactions were then cleaned using the QIAquick PCR Purification Kit (Qiagen, Cat # 28104). To do so, the cleaned fragments were first concentrated in 18 μL of water via elution. Then, the eluate was added to 2 μL of 1X E-Gel Sample Loading Buffer (Thermo Fisher Scientific, cat # 10482055) and the resulting 20 μL loaded onto a well on a 2% Agarose E-gel (Thermo Fisher Scientific, Cat # G401002). The E-gel was then loaded onto the E-Gel Power Snap Electrophoresis Device (Thermo Fisher Scientific, Cat # G8100) and electrophoresed for 13 mins on the 1-2% agarose gel setting. Finally, a band corresponding to the size of the desired fragment was excised from the agarose gel and purified using the Qiagen min-elute gel extraction kit (Qiagen, Cat # 28604). Next, the purified DNA fragments were eluted in 14 μL water and quantified on a Nanodrop spectrophotometer. When the fragments were to be used for targeted sequencing, the purified fragments were quantified using a Qubit 3.0 fluorometer (Thermo Fisher Scientific, cat # Q33216) with the dsDNA broad range kit (Thermo Fisher Scientific, cat # Q32850).

### Poly(A)-enriched RNA sequencing

To prepare poly(A)-enriched RNA sequencing libraries, we processed 500 ng of total RNA using the TruSeq Stranded mRNA Library Prep kit (Illumina, Cat # 20020595; Indexes: IDT for Illumina TruSeq RNA UD Indexes, cat # 20020591). The resulting libraries were then analyzed on a TapeStation (Agilent) and quantified via a Qubit 3.0 fluorometer. Equimolar amounts of the resulting sequencing libraries were combined to make a 4 nM pool. The pools were then sequenced on a Novaseq 6000 instrument (Illumina) in single-end 100 bp format at the Institute for Genomic Medicine at UCSD or the La Jolla Institute for Immunology.

### Standard sequencing library normalization

For standard normalization, we calculated the concentration in nM for each library using the equation: concentration in nM = [(concentration in ng/μL) ÷ (660 g/mol × average library size in bp)] × 10^6^. The average library size was obtained via analysis of 1 μL of the library on the Agilent TapeStation 2200 (Agilent, Cat # G2964AA) using D1000 screen tape (Agilent, Cat # 5067-5583). The library concentration in ng/μL was determined by analysis of 1 μL of the library on the Qubit 3.0 fluorometer (Thermo Fisher Scientific, cat # Q33216) using the dsDNA broad range kit (Thermo Fisher Scientific, cat # Q32850).

### Next-generation sequencing

NovaSeq sequencing was carried out on the Novaseq 6000 at the La Jolla Institute for Immunology (LJI) or the UC San Diego Health Sciences Institute for Genomic Medicine (IGM) Genomics Center. Further, MiSeq sequencing was done at the Stem Cell Genomics and Microscopy Core at the Sanford Consortium for Regenerative Medicine (SCRM).

### RPS2-BE editing +/- Torin-1 treatment

For mTOR perturbation experiments, cells were transiently transfected with the RPS2-BE fusions as described above (see transient transfection section), and expression of the constructs was induced via the addition of doxycycline at a final concentration of 2 μg mL^−1^ and incubated at 37 °C for 24 hrs. After, Torin-1 (Cell Signaling, Cat # 14379) dissolved in dimethyl sulfoxide (DMSO) was added to pre-warmed DMEM +10% FBS, adding the media to the cultured cells yielded a final concentration of 100 nM. A set of cells were treated with a DMSO vehicle lacking Torin-1 to serve as a control. The cells were then incubated at 37 °C for 48 hrs., after which the media was aspirated, and cells were resuspended in 300 μL of TRIzol followed by RNA extraction, library prep, and sequencing (see above). To ensure the reliability of the results, the experiments were carried out on three replicates.

### Dual editing

#### The reporter-based dual editing system

The 12X MS2-SL plasmid was digested with AfeI (New England Biolabs, Cat # R0652S) or FastDigest Eco47III (Thermo Fisher Scientific, Cat # FD0324) and PmeI (New England Biolabs, Cat # R0560S) to swap in different MS2 and PP7 stem-loop distributions for the reporter-based dual-editing system. Gene blocks (IDT) coding one of the three distinct MS2 and PP7 distributions considered were cloned into the purified backbone via Gibson assembly (See detailed Gibson cloning procedure above). Further, the protein component was generated by digesting the plasmid encoding the MCP-8e-HA-P2A-mRuby construct with NheI (Thermo Fisher Scientific, Cat # FD0974) and AfeI (New England Biolabs, Cat # R0652S) or FastDigest Eco47III (Thermo Fisher Scientific, Cat # FD0324) to remove the MCP coding sequence. A gene block encoding the PP7- CP^57^ was then Gibson cloned into the purified backbone to generate PP7-CP-8e-HA-P2A-mRuby. After the newly-cloned construct was digested with AfeI (New England Biolabs, Cat # R06 Thuronyi 2S) or FastDigest Eco47III (Thermo Fisher Scientific, Cat # FD0324) and BshTI (AgeI) (Thermo Fisher Scientific, Cat # FD1464) to remove the 8e-HA-P2A-mRuby-coding sequences and PCR products encoding 8e-HA-P2A or 7.10 (V82G)-HA-P2A were cloned into the purified backbone together with a PCR product encoding the blue fluorescent protein^58^ (BFP) coding sequence. The reaction yielded PP7-CP-8e-HA-P2A-BFP and PP7-CP-7.10 (V82G)-HA-P2A- BFP. The resulting PP7-A-to-I editor-encoding plasmids were then individually co-transfected into HEK293XT cells with each of the MS2- and PP7-stem-loop-bearing reporters (one reporter at a time) and a plasmid encoding the MCP-APOBEC1-HA-P2A-mRuby C-to-U-editing construct. Experiments in which each editing fusion was co-transfected with the reporter without a second editor were done to help determine the specificity of the coat proteins for their given stem-loops. Experiments in which the reporter was transfected without an editor were done to help account for editing that may arise from endogenous enzymes or sequencing errors. All experiments were done in duplicate to ensure the reproducibility of the results.

### Data Processing

#### MS2 Loop Specificity Data

Cutadapt was used to remove adapter sequences from original FASTQ files, after which bwa mem was used to align sequences to a FASTA file containing a single entry, representing the twelve-loop construct sequence. Finally, Pysamstats was used to generate count tables of base counts at each position along the construct, which was processed using the R statistical software ggplot2 to visualize editing levels as a histogram. The histogram displays Green (C-to-U), orange (A-to-I), and black “No Edit” bars. To make it easier to see the C-to-U and A-to-I edit bars, only the top line of the black “No Edit” bars was kept, which produced the “No Edit” line.

### MS2 Loop On/Off-Target Data (signal-to-noise)

Cutadapt was used to remove adapter sequences from the original FASTQ files, after which the resulting FASTQ files were aligned to the GRCh38 human reference genome using STAR 2.7.6a. The STAR default settings were used, except for *--outSAMtype BAM SortedByCoordinate* and *--outSAMunmapped Within*. Unmapped and mapped reads were extracted from the resulting bam, using the commands *samtools view -f4* and *samtools view -F4*, respectively, into separate bams. Reads from the .bam file with unmapped reads were converted back to a FASTQ file, which was subsequently aligned to a FASTA file containing a single entry representing the twelve-loop construct sequence, again using the STAR 2.7.6a alignment software. SAILOR was then run twice (once for A-to-I edit detection and once for C-to-T edit detection) on all reference-mapped and construct-mapped .bam files, using either the reference genome or the twelve-loop construct FASTA file as a reference, respectively. Finally, SAILOR edit counts were loaded into a Jupyter notebook. On/off-target rates were then calculated by dividing, for each sample, the edit count on reads mapped to the reporter construct (on-target) by the edit count on reads mapped to the genome (off-target). This ensured, for example, that enzymes that have high on-target but also high off-target have a lower signal-to-noise ratio than enzymes with low on-target but much lower off-target.

### RBFOX2-rBE data

Cutadapt was used to trim adapter sequences from reads in all original FASTQ files, after which the resulting trimmed reads were aligned to the GRCh38 human reference genome using STAR 2.7.6 with default settings except for *--outSAMtype BAM SortedByCoordinate* and *-- outSAMunmapped Within.* SAILOR was run for each bam file, with parameters variably set for A-to-I detection or C-to-T detection, depending on the known editing modality of each enzyme being tested. C-to-T and A-to-I SAILOR output files were combined for A2dd (R), which is known to exhibit both editing modalities. For each replicate of each enzyme, edit counts were summed for 30-bp bins across regions of the transcriptome exhibiting edits. Using a background rate calculated as the mean editing fraction (fraction of editable Cs or As, respectively, edited per bin) across all bins, a Poisson test was conducted for each bin to test for significantly elevated editing levels. Benjamini-Hochberg false discovery rate correction was used to adjust p-values, and only bins with adjusted p-values below 0.1 were retained. After filtering bins, the remaining bins within 15 bp of each other were merged to form clusters. For each enzyme, peak coordinates were intersected between all three replicates, and only clusters present in all three replicates were retained. The clusters were loaded into Jupyter notebooks, where motif presence, and RBFOX2- APOBEC1 eCLIP overlap were calculated using custom scripts.

### RPS2 +/- Torin1 data

Cutadapt was used to trim adapter sequences from reads in all original FASTQ files, after which the resulting trimmed reads were aligned to the GRCh38 human reference genome using STAR 2.7.6 with default settings except for *--outSAMtype BAM SortedByCoordinate* and *-- outSAMunmapped Within.* SAILOR was run for each bam file, with parameters variably set for A-to-I detection or C-to-T detection, depending on the known editing modality of each enzyme being tested. C-to-T and A-to-I SAILOR output files were combined for A2dd (R), which is known to exhibit both editing modalities. The subread featurecounts software was used to obtain read counts for genes in all samples, and then edits per read (EPR) were calculated on a per-gene basis using the output of featurecounts and the outputs of SAILOR. EPR data was loaded into Jupyter notebooks, where relative decreases of editing under the influence of Torin-1 were calculated for each sample.

### Reporter-based dual editing sequencing data

Reference FASTA and GTF formatted files were prepared for each designed reporter. Adapters were trimmed from short reads using cutadapt. A Burrows-Wheeler Alignment Tool (BWA) (version 0.7.17) index was then built for each reporter and BWA mem was used with default parameters to align reads to the respective reporter sequence. Edits were identified with pysamstats (version 1.1.2) using parameters: *--min-baseq 20* and *--type variation_strand.* Plots to visualize sites containing A-to-I (G) and C-to-T edits were created using a custom R script.

### Reproducibility

To ensure reproducibility, several experiments were conducted using two replicates. These experiments included MS2 reporter assays, on-to-off-target analyses, and integrated C-to-U and A-to-I (MCP and PP7) experiments. The data from the reporter experiments were analyzed independently per replicate, while the edit values for both replicates were combined for the on-to-off-target analyses before comparing reporter (on-target) versus transcriptome (off-target).

The RBFOX2-rBE and RPS2-rBE experiments were performed on three replicates. For the RBFOX2-rBE data, only FLARE edit clusters that were consistent across all three replicates were considered for analysis. Additionally, clusters that were detected consistently across three rBE-only experiments were treated as noise and subtracted from the RBFOX2 data. For the RPS2-rBE experiments, only edits on mRNAs edited across three independent replicates were considered for analyses.

## Acknowledgements

H.C.M.M. was supported by a UCSD Institutional Research and Academic Career Development Award (IRACDA). G.D.L. was supported by an IRACDA Summer Undergraduate Research Fellowship (SURF). G.W.Y. is supported as an Allen Distinguished Investigator, a Paul G. Allen Frontier Group advised grant to the Paul G. Allen Family Foundation. G.W.Y. is supported by NIH Grants U41HG009889, RF1MH126719, R01NS103172, R01HG004659 and R01HG011864.

RMK is supported by NIH Grant R01-HG-010646. A.C.K. was partially supported by the NIH through grant no. 1R35GM138317 (to A.C.K). B.L.R. was supported by the Chemistry-Biology Interface Training Program, NIH Grant T32 GM112584.

## Contributions

Conceptualization: H.C.M.M. and G.W.Y.; methodology: H.C.M.M..; investigation: H.C.M.M., T.Y., K.L.J., J.R.M., G.D.L., A.T.D., S.S.P., S.M.B., and B.L.R.; formal analysis: E.K., P.J., E.A.B., and H.C.M.M.; writing of original draft: H.C.M.M. and G.W.Y.; writing of review and editing: G.W.Y., H.C.M.M., E.K., P.J., R.M.K., and A.C.K.; funding acquisition: H.C.M.M. and G.W.Y.; supervision: G.W.Y.

## Biological materials

The plasmids constructed for this study will be deposited to Addgene for distribution under a Uniform Biological Material Transfer Agreement (UBMTA).

## Data availability

Raw and assembled sequencing data from this study will be deposited in NCBI’s Gene Expression Omnibus (GEO). Accession codes for all relevant data will be provided before publishing.

## Code availability

Source code and analysis scripts for edit quantification will be available as Supplementary Software before publishing.

## Supplementary Figures

**Supplementary Figure 1:**
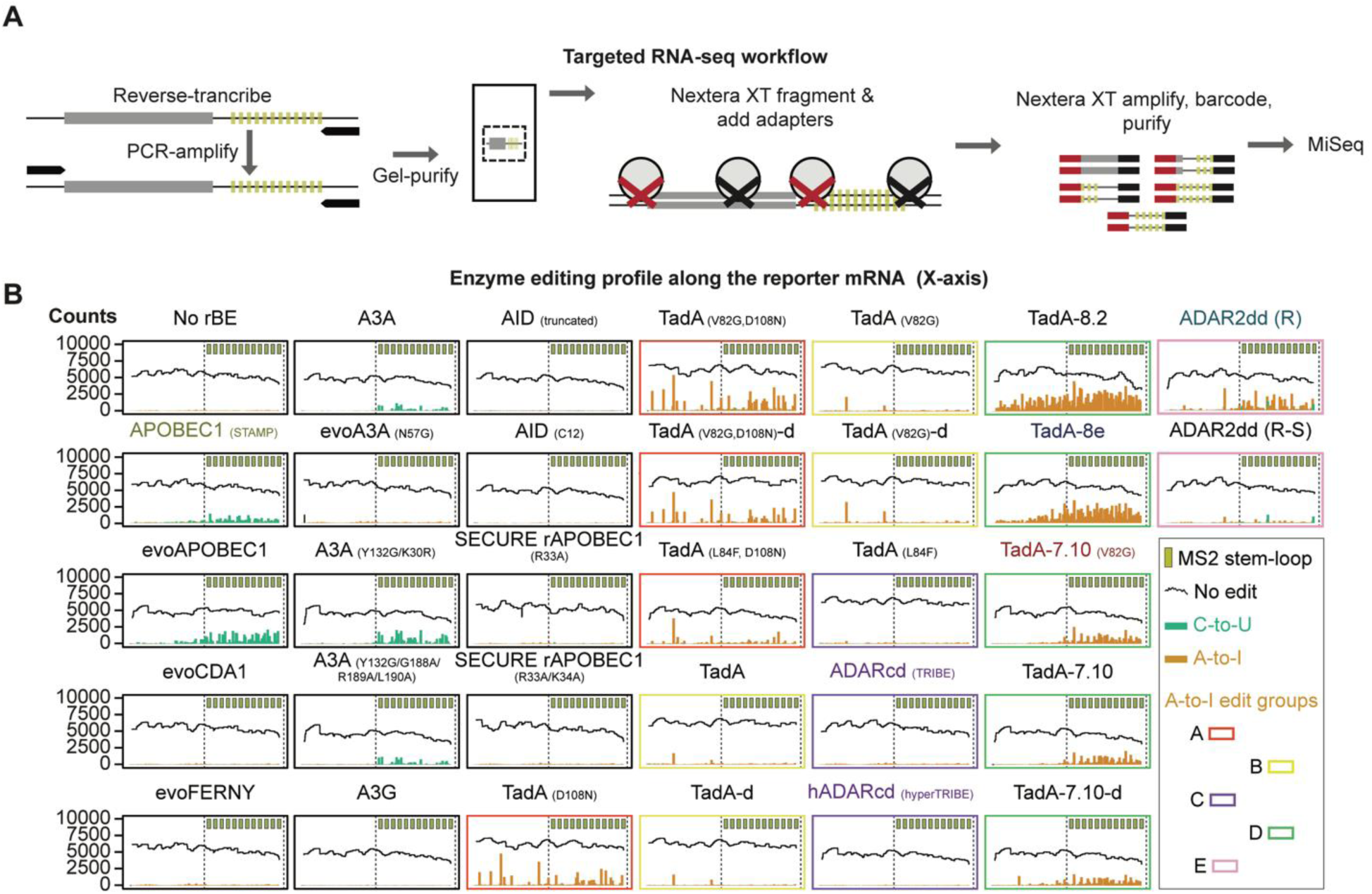
Detecting edits on the reporter mRNA. **A)** Targeted RNA-seq strategy to detect edits along the reporter mRNA sequence. **B)** The number of times a base was called (y-axis) at each position along the twelve-MS2 stem- loop reporter construct (as in Figure 1). The fraction of each position in the reporter that contained either a cytosine (C) but a uridine was detected (C-to-U) or an adenosine (A) but an inosine was detected (read as guanosine, A-to-I) are denoted by green and orange bars, respectively. Positions where the called base matches the reporter sequence (no edit) are indicated by black bars.

**Supplementary Figure 2:**
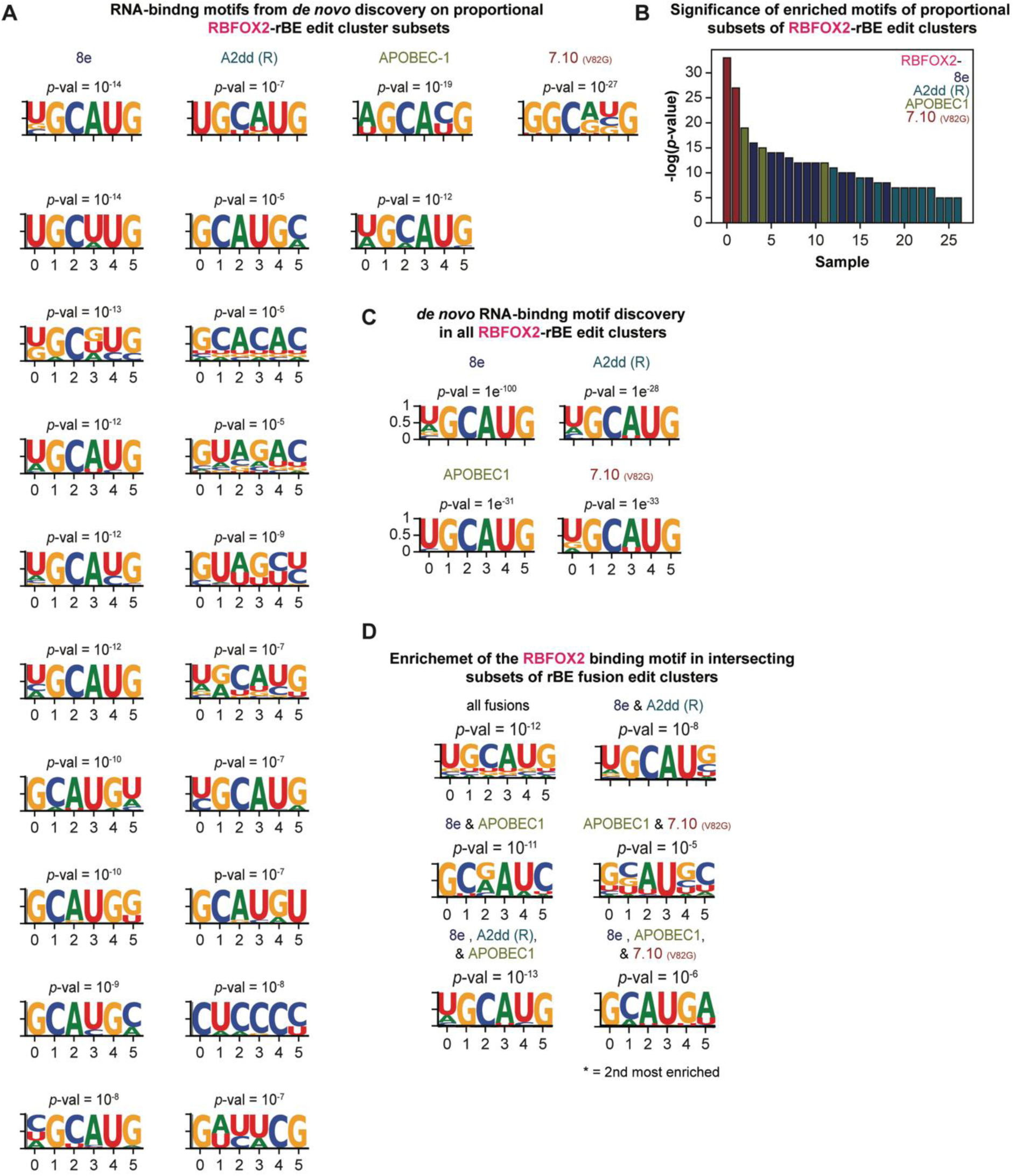
HOMER sequence motif enrichment analyses detect enrichment of the expected RBFOX2 sequence motif. **A)** HOMER motif enriched in an equal number of edit clusters sampled from each RBFOX2-rBE fusion set. 736 edit clusters were repeatedly selected from each cluster set at random and without replacement until no further sets of 736 clusters could be chosen. The most enriched sequence motif produced by a selection of 736 clusters from each fusion is shown in figure 2E, and the motifs for selections that yielded lower p-values are shown here in order of most (top) to least (bottom) significant. **B)** The negative logarithm-transformed p-values (-log *p*-value) associated with the HOMER motifs listed in Supplementary Figure 2A. **C)** HOMER detects the RBFOX2 motif among each set of RBFOX2-rBE edit clusters. **D)** The RBFOX2 binding motifs are the most or second most (*) enriched HOMER motif among each set of intersecting peaks (see also Figure 2I).

**Supplementary Figure 3.**
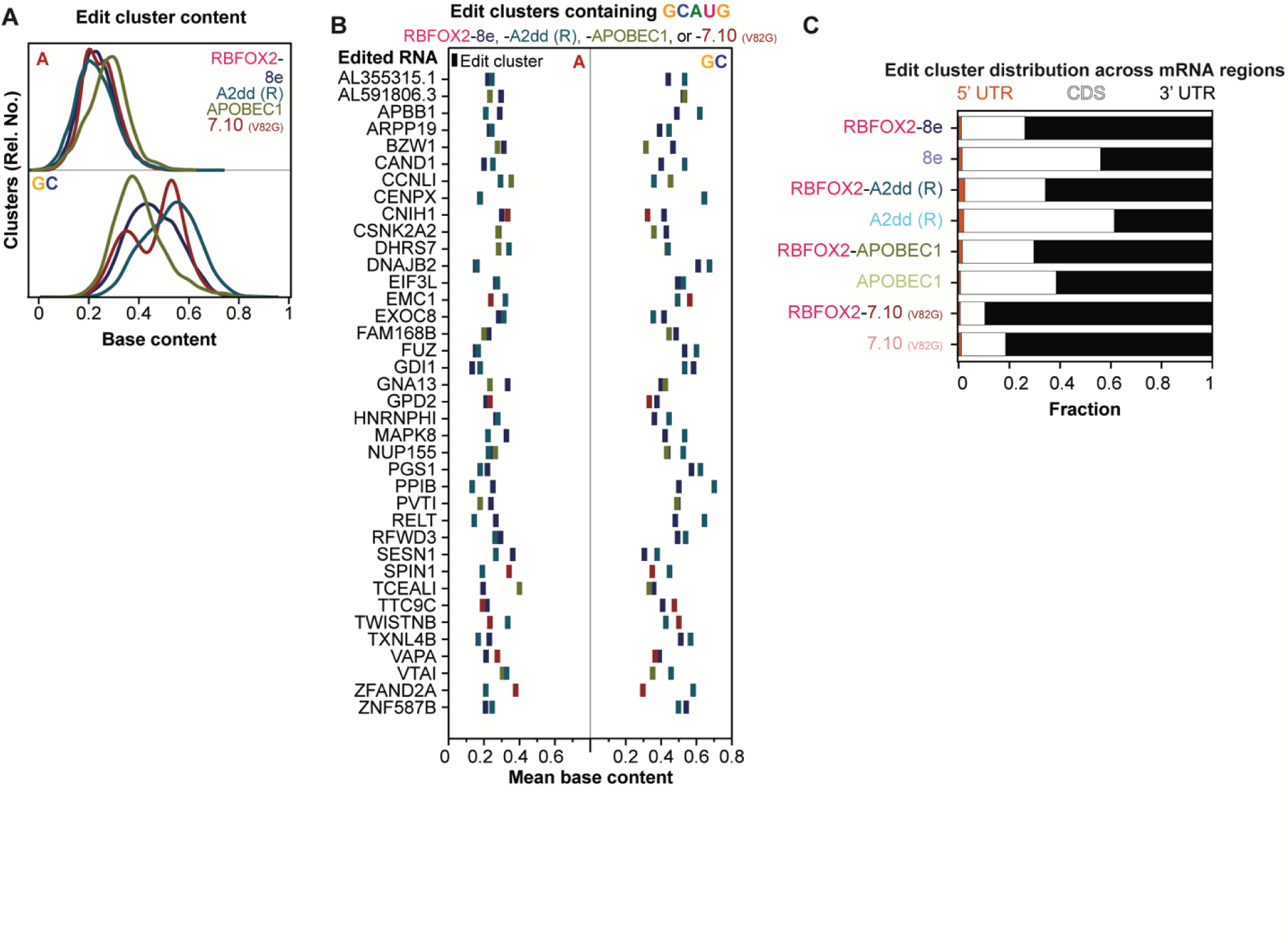
**A)** A density plot of the overall peak adenosine (A, top) and guanosine-cytosine (GC, bottom) content for RBFOX2-rBE fusions before subtraction of the edit clusters produced by the corresponding rBE-only construct (see Figure 3E for density plots of the remaining RBFOX2-rBE and all rBE only edit clusters). **B)** On genes with two GCAUG core motif-containing clusters edited by two different enzymes, the combination of flanking window base contents and enzyme editing context specificities depicted in panel E dictate which region is more likely to be edited by each RBFOX2-rBE fusion. **C)** Analyses of where on the mRNA region the RBFOX2-rBE and rBE only edit clusters map to. Bars demonstrate the fraction of edit clusters that map to the 5’ (red) or 3’ (black) untranslated region (UTR), and those mapping to the coding sequence (CDS, white).

**Supplementary Figure 4:**
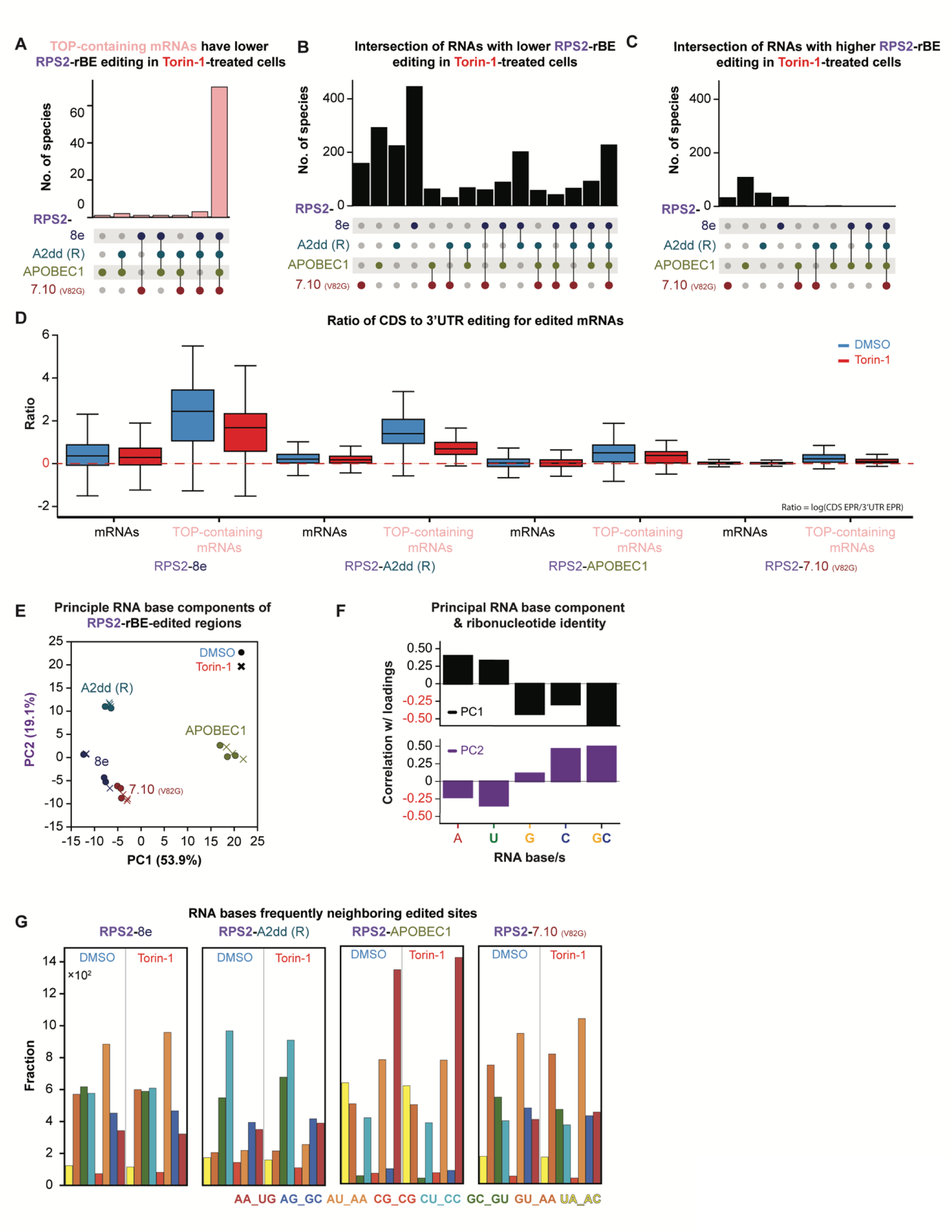
RNAs that exhibited significant RPS2-rBE edits per read (EPR) differences in cells grown under translation (Torin-1-treated) and uninhibited conditions (DMSO-treated) (*see methods*). **A-C)** The intersection between RBFOX2-rBE constructs for **A)** TOP-containing (pink bars) and **B)** all edited poly(A)+ RNAs (black) that experienced a lower EPR in Torin-1-treated cells relative to the DMSO vehicle-treated cells, and **C)** all poly(A)+ RNAs (black) with an increased EPR. **D)** The ratio of RPS2-rBE edits between mRNA coding sequences (CDSs) and 3’ untranslated regions (3’UTRs) for TOP-containing (pink letters) and all poly(A)+ RNAs (black letters). The plot boxes are colored based on whether the cells were grown in uninhibited (DMSO, blue) or translation-inhibiting (Torin-1-treated, red) conditions. **E)** PCA analysis of bases flanking edited sites. **F)** Contribution of each RNA base (A, U, G, or C) or combination (GC) to principal components PC1 (black) and PC2 (purple). **G)** The top pairs of RNA bases flanking sites edited by each RPS2-rBE fusion. The bars denote the fraction of the edited sites within a given context, and the bars are colored in agreement with the corresponding sequence context listed at the bottom.

**Supplementary Figure 5:**
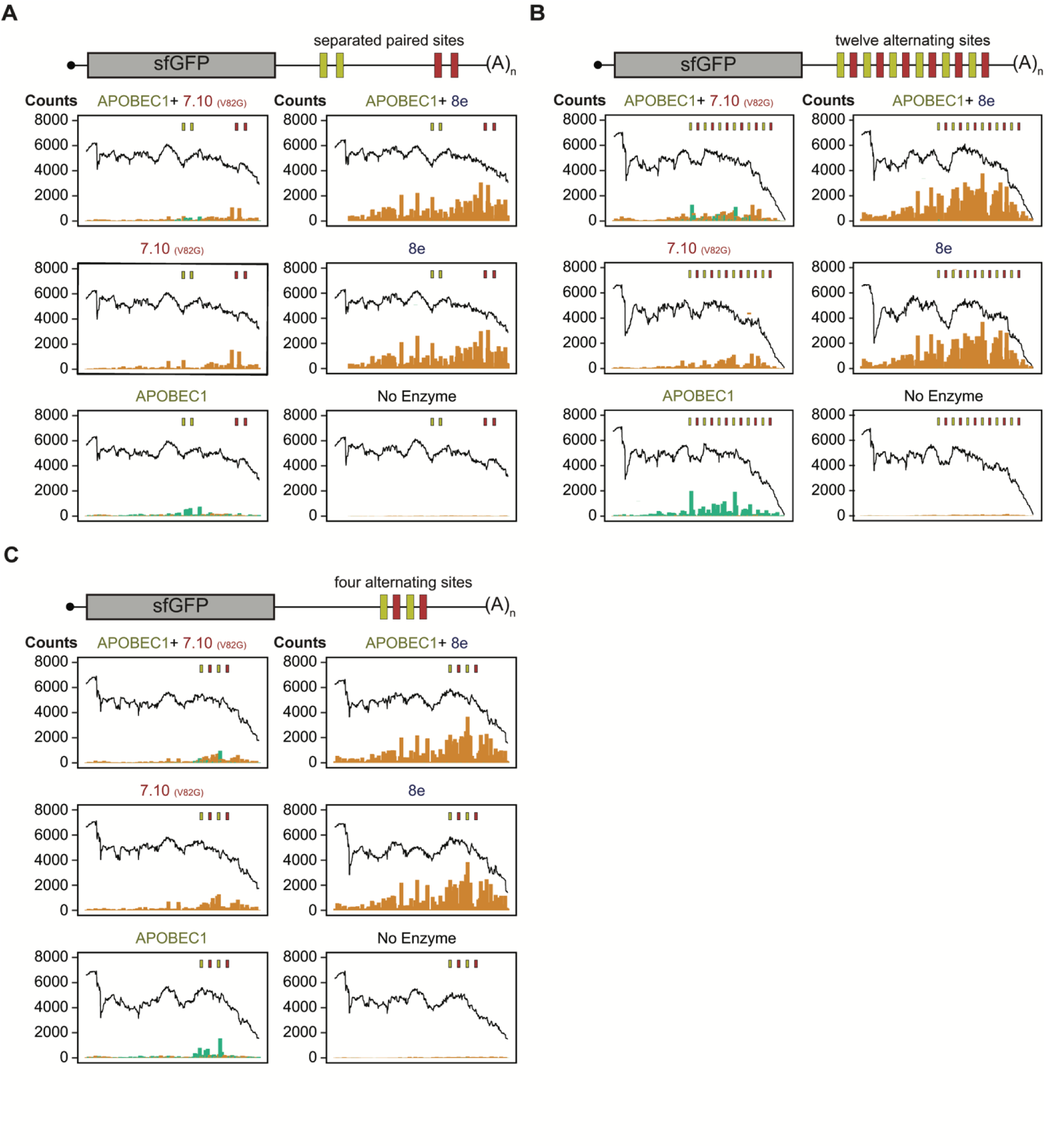
The combinatorial editing profiles for replicates of experiments shown in Figure 5. The combinatorial editing profiles for MCP-APOBEC1 together with either PP7-CP-8e or PP7- CP-7.10 (V82G) when they are co-expressed in HEKs with a reporter mRNA bearing MS2 and PP7 stem-loops in **A)** split pairs or **B)** twelve and **C)** four alternating sites. The fraction of total covered bases at each position along each reporter exhibiting either C-to-U (green) or A-to-I (orange) are denoted as bars. In contrast, the fraction of reads with no edits is denoted by a black line.

## References

1 Gebauer, F., Schwarzl, T., Valcárcel, J. & Hentze, M. W. RNA-binding proteins in human genetic disease. Nature Reviews Genetics 22, 185–198 (2021). 10.1038/s41576-020-00302-y

2 Ingolia, N. T., Ghaemmaghami, S., Newman, J. R. S. & Weissman, J. S. Genome-Wide Analysis in Vivo of Translation with Nucleotide Resolution Using Ribosome Profiling. Science 324, 218–223 (2009). 10.1126/science.1168978

3 Van Nostrand, E. L. et al. Robust transcriptome-wide discovery of RNA-binding protein binding sites with enhanced CLIP (eCLIP). Nature Methods 13, 508–514 (2016). 10.1038/nmeth.3810

4 McMahon, Aoife C. et al. TRIBE: Hijacking an RNA-Editing Enzyme to Identify Cell-Specific Targets of RNA-Binding Proteins. Cell 165, 742–753 (2016). 10.1016/j.cell.2016.03.007

5 Brannan, K. W. et al. Robust single-cell discovery of RNA targets of RNA-binding proteins and ribosomes. Nature Methods 18, 507–519 (2021). 10.1038/s41592-021-01128-0

6 Kofman, E., Yee, B. A., Medina-Munoz, H. C. & Yeo, G. W. FLARE: a fast and flexible workflow for identifying RNA editing foci. BMC Bioinformatics (2023). DOI: 10.1186/s12859-023-05452-4

7 Rosenberg, B. R., Hamilton, C. E., Mwangi, M. M., Dewell, S. & Papavasiliou, F. N. Transcriptome-wide sequencing reveals numerous APOBEC1 mRNA-editing targets in transcript 3’ UTRs. Nat Struct Mol Biol 18, 230–236 (2011). 10.1038/nsmb.1975

8. 8 Thuronyi, B. W., et al. Continuous evolution of base editors with expanded target compatibility and improved activity. Nature Biotechnology 37, 1070–1079 (2019). 10.1038/s41587-019-0193-0

9 Macbeth, M. R. et al. Inositol Hexakisphosphate Is Bound in the ADAR2 Core and Required for RNA Editing. Science 309, 1534–1539 (2005). 10.1126/science.1113150

10 Eggington, J. M., Greene, T. & Bass, B. L. Predicting sites of ADAR editing in double-stranded RNA. Nature Communications 2, 319 (2011). 10.1038/ncomms1324

11 Montiel-Gonzalez, M. F., Vallecillo-Viejo, I., Yudowski, G. A. & Rosenthal, J. J. C. Correction of mutations within the cystic fibrosis transmembrane conductance regulator by site-directed RNA editing. Proceedings of the National Academy of Sciences 110, 18285–18290 (2013). 10.1073/pnas.1306243110

12 Vogel, P., Schneider, M. F., Wettengel, J. & Stafforst, T. Improving Site-Directed RNA Editing In Vitro and in Cell Culture by Chemical Modification of the GuideRNA. Angewandte Chemie International Edition 53, 6267–6271 (2014). 10.1002/anie.201402634

13 Vogel, P. & Stafforst, T. Site-Directed RNA Editing with Antagomir Deaminases — A Tool to Study Protein and RNA Function. ChemMedChem 9, 2021–2025 (2014). 10.1002/cmdc.201402139

14 Phelps, C. B. & Brand, A. H. Ectopic Gene Expression inDrosophilaUsing GAL4 System. Methods 14, 367–379 (1998). 10.1006/meth.1998.0592

15 Xu, W., Rahman, R. & Rosbash, M. Mechanistic implications of enhanced editing by a HyperTRIBE RNA-binding protein. Rna 24, 173–182 (2018). 10.1261/rna.064691.117

16 Tutucci, E. et al. An improved MS2 system for accurate reporting of the mRNA life cycle. Nature Methods 15, 81–89 (2018). 10.1038/nmeth.4502

17 Abudayyeh, O. O. et al. A cytosine deaminase for programmable single-base RNA editing. Science 365, 382–386 (2019). 10.1126/science.aax7063

18 Grünewald, J., et al. CRISPR DNA base editors with reduced RNA off-target and self-editing activities. Nature Biotechnology 37, 1041–1048 (2019). 10.1038/s41587-019-0236-6

19 Komor, A. C., Kim, Y. B., Packer, M. S., Zuris, J. A. & Liu, D. R. Programmable editing of a target base in genomic DNA without double-stranded DNA cleavage. Nature 533, 420–424 (2016). 10.1038/nature17946

20 Nishida, K. et al. Targeted nucleotide editing using hybrid prokaryotic and vertebrate adaptive immune systems. Science 353, aaf8729 (2016). doi:10.1126/science.aaf8729

21 Gaudelli, N. M. et al. Programmable base editing of A•T to G•C in genomic DNA without DNA cleavage. Nature 551, 464–471 (2017). 10.1038/nature24644

22 Koblan, L. W., et al. Improving cytidine and adenine base editors by expression optimization and ancestral reconstruction. Nature Biotechnology 36, 843–846 (2018). 10.1038/nbt.4172

23. 23 Gehrke, J. M., et al. An APOBEC3A-Cas9 base editor with minimized bystander and off-target activities. Nature Biotechnology 36, 977–982 (2018). 10.1038/nbt.4199

24 Grünewald, J., et al. A dual-deaminase CRISPR base editor enables concurrent adenine and cytosine editing. Nature Biotechnology 38, 861–864 (2020). 10.1038/s41587-020-0535-y

25 Gaudelli, N. M., et al. Directed evolution of adenine base editors with increased activity and therapeutic application. Nature Biotechnology 38, 892–900 (2020). 10.1038/s41587-020-0491-6

26 Richter, M. F., et al. Phage-assisted evolution of an adenine base editor with improved Cas domain compatibility and activity. Nature Biotechnology 38, 883–891 (2020). 10.1038/s41587-020-0453-z

27 Zuo, E., et al. A rationally engineered cytosine base editor retains high on-target activity while reducing both DNA and RNA off-target effects. Nature Methods 17, 600–604 (2020). 10.1038/s41592-020-0832-x

28 Kurt, I. C., et al. CRISPR C-to-G base editors for inducing targeted DNA transversions in human cells. Nature Biotechnology 39, 41–46 (2021). 10.1038/s41587-020-0609-x

29 Barka, A. et al. The Base-Editing Enzyme APOBEC3A Catalyzes Cytosine Deamination in RNA with Low Proficiency and High Selectivity. ACS Chemical Biology 17, 629–636 (2022). 10.1021/acschembio.1c00919

30 Tang, G. et al. Creating RNA Specific C-to-U Editase from APOBEC3A by Separation of Its Activities on DNA and RNA Substrates. ACS Synthetic Biology 10, 1106–1115 (2021). 10.1021/acssynbio.0c00627

31 Kohli, R. M. et al. Local sequence targeting in the AID/APOBEC family differentially impacts retroviral restriction and antibody diversification. J Biol Chem 285, 40956–40964 (2010). 10.1074/jbc.M110.177402

32 Kohli, R. M. et al. A portable hot spot recognition loop transfers sequence preferences from APOBEC family members to activation-induced cytidine deaminase. J Biol Chem 284, 22898–22904 (2009). 10.1074/jbc.M109.025536

33. 33 Berríos, K. N., et al. Controllable genome editing with split-engineered base editors. Nat Chem Biol 17, 1262–1270 (2021). 10.1038/s41589-021-00880-w

34. 34 Rallapalli, K. L., Komor, A. C. & Paesani, F. Computer simulations explain mutation-induced effects on the DNA editing by adenine base editors. Sci Adv 6, eaaz2309 (2020). 10.1126/sciadv.aaz2309

35 Johansson, H. E. et al. A thermodynamic analysis of the sequence-specific binding of RNA by bacteriophage MS2 coat protein. Proc Natl Acad Sci U S A 95, 9244–9249 (1998). 10.1073/pnas.95.16.9244

36 Jin, Y. et al. A vertebrate RNA-binding protein Fox-1 regulates tissue-specific splicing via the pentanucleotide GCAUG. The EMBO Journal 22, 905–912 (2003). 10.1093/emboj/cdg089

37. 37 Washburn, M. C., et al. The dsRBP and inactive editor ADR-1 utilizes dsRNA binding to regulate A-to-I RNA editing across the C. elegans transcriptome. Cell Rep 6, 599–607 (2014). 10.1016/j.celrep.2014.01.011

38 Heinz, S. et al. Simple Combinations of Lineage-Determining Transcription Factors Prime *cis*-Regulatory Elements Required for Macrophage and B Cell Identities. Molecular Cell 38, 576–589 (2010). 10.1016/j.molcel.2010.05.004

39 Yeo, G. W. et al. An RNA code for the FOX2 splicing regulator revealed by mapping RNA-protein interactions in stem cells. Nat Struct Mol Biol 16, 130–137 (2009). 10.1038/nsmb.1545

40 Zhou, D., Couture, S., Scott, M. S. & Abou Elela, S. RBFOX2 alters splicing outcome in distinct binding modes with multiple protein partners. Nucleic Acids Research 49, 8370–8383 (2021). 10.1093/nar/gkab595

41 Rahman, R., Xu, W., Jin, H. & Rosbash, M. Identification of RNA-binding protein targets with HyperTRIBE. Nature Protocols 13, 1829–1849 (2018). 10.1038/s41596-018-0020-y

42 Blanc, V. et al. Genome-wide identification and functional analysis of Apobec-1-mediated C-to-U RNA editing in mouse small intestine and liver. Genome Biology 15, R79 (2014). 10.1186/gb-2014-15-6-r79

43 McDaniel, Y. Z. et al. Deamination hotspots among APOBEC3 family members are defined by both target site sequence context and ssDNA secondary structure. Nucleic Acids Research 48, 1353–1371 (2020). 10.1093/nar/gkz1164

44 Jia, J.-J. et al. mTORC1 promotes TOP mRNA translation through site-specific phosphorylation of LARP1. Nucleic Acids Research 49, 3461–3489 (2021). 10.1093/nar/gkaa1239

45 Flamand, M. N., Ke, K., Tamming, R. & Meyer, K. D. Single-molecule identification of the target RNAs of different RNA binding proteins simultaneously in cells. Genes & Development (2022). 10.1101/gad.349983.122

46 Chen, L.-L., Yang, L. & Carmichael, G. Molecular basis for an attenuated cytoplasmic dsRNA response in human embryonic stem cells. Cell Cycle 9, 3552–3564 (2010). 10.4161/cc.9.17.12792

47 Yizhu, L. et al. Single molecule co-occupancy of RNA-binding proteins with an evolved RNA deaminase. bioRxiv, 2022.2009.2006.506853 (2022). 10.1101/2022.09.06.506853

48 Katharine, C. A., Corrie, R. & Michael, R. Comparison of TRIBE and STAMP for identifying targets of RNA binding proteins in human and *Drosophila* cells. bioRxiv, 2023.2002.2003.527025 (2023). 10.1101/2023.02.03.527025

49 Medina-Muñoz, H. C., Lapointe, C. P., Porter, D. F. & Wickens, M. Records of RNA localization through covalent tagging. bioRxiv, 785816 (2019). 10.1101/785816

50 Medina-Munoz, H. C., Lapointe, C. P., Porter, D. F. & Wickens, M. Records of RNA locations in living yeast revealed through covalent marks. Proc Natl Acad Sci U S A 117, 23539–23547 (2020). 10.1073/pnas.1921408117

51 Jan, C. H., Williams, C. C. & Weissman, J. S. Principles of ER cotranslational translocation revealed by proximity-specific ribosome profiling. Science 346, 1257521 (2014). 10.1126/science.1257521

52 Williams, C. C., Jan, C. H. & Weissman, J. S. Targeting and plasticity of mitochondrial proteins revealed by proximity-specific ribosome profiling. Science 346, 748–751 (2014). 10.1126/science.1257522

53 Fazal, F. M. et al. Atlas of Subcellular RNA Localization Revealed by APEX-Seq. Cell 178, 473–490.e426 (2019). 10.1016/j.cell.2019.05.027

54 Pédelacq, J.-D., Cabantous, S., Tran, T., Terwilliger, T. C. & Waldo, G. S. Engineering and characterization of a superfolder green fluorescent protein. Nature Biotechnology 24, 79–88 (2006). 10.1038/nbt1172

55 Gibson, D. G. et al. Enzymatic assembly of DNA molecules up to several hundred kilobases. Nature Methods 6, 343–345 (2009). 10.1038/nmeth.1318

56 Hartley, J. L., Temple, G. F. & Brasch, M. A. DNA cloning using in vitro site-specific recombination. Genome Res 10, 1788–1795 (2000). 10.1101/gr.143000

57 Dhaese, P., Lenaerts, A., Gielen, J. & Van Montagu, M. Complete amino acid sequence of the coat protein of the Pseudomonas aeruginosa RNA bacteriophage PP7. Biochem Biophys Res Commun 94, 1394–1400 (1980). 10.1016/0006-291x(80)90574-4

58 Wu, B., Eliscovich, C., Yoon, Y. J. & Singer, R. H. Translation dynamics of single mRNAs in live cells and neurons. Science 352, 1430–1435 (2016). 10.1126/science.aaf1084

